# “Single-Nuclei Analysis of the Unfolded Protein Response (SNUPR) reveals bortezomib resistance mechanisms in Multiple Myeloma”

**DOI:** 10.1101/2024.10.22.617161

**Authors:** Julien P. Gigan, Paulina Garcia-Gonzalez, Lou Galliot, Alexandre Reynaud, Yoan Ghaffar, Felipe Flores-Santibáñez, Eve Seillier, Rosario Lavignolle-Heguy, Daniela Barros Dos Santos, Sharon Fischaux, Alexis Combes, Miwako Narita, Evelina Gatti, Beatrice Nal, Stéphane Rocchi, Jerome Moreaux, Philippe Pierre, Rafael J. Argüello

## Abstract

The unfolded protein response (UPR) is a key stress resistance pathway that has become a key potential target for improving the efficacy of cancer chemotherapy. The UPR involves the activation of three ER-resident stress sensors: PERK, IRE-1 and ATF6 with different signalling outcomes leading to cell death or survival. These cell-fate decisions are difficult to predict and are the result of the complex interaction of PERK, IRE-1 and ATF6 downstream events that have differences in their dynamics and their interplay. These characteristics of the UPR are still poorly defined due to lack of methods to monitor their activation simultaneously at single-cell level. We developed SNUPR (Single Nuclei analysis of the Unfolded Protein Response), an accessible technique that allows the profiling of the three UPR branches in nuclear suspensions by flow cytometry, and applied it to study UPR dynamics in a cancer-specific context. By performing transcriptomic analysis, we found that ER-stress sensor specific gene signatures correlate with patient survival in several blood malignancies, and by using SNUPR, we detected high heterogeneity during UPR activation *in vitro* in different human cancer cell lines, which could not be have been predicted by the level of expression of the sensors. Our SNUPR analyses further indicate that this heterogeneity is explained by variations in the intensity and duration of ER stress-induced protein synthesis inhibition via PERK, acting as upstream regulator of both the IRE-1/XBP1 and ATF6 dependent transcriptional programs. We extend the relevance of these observations by demonstrating that IRE-1/XBP1s pathway plays a critical role in bortezomib resistance of multiple myeloma cells and patients. We present here SNUPR, that can be used to monitor UPR dynamics with single-cell resolution and identified clinical contexts in which targeting a specific UPR branch could be detrimental or help circumventing chemotherapy resistance.

**One Sentence Summary:** SNUPR method enable single-cell UPR profiling and reveals the role of IRE-1 axis in predicting bortezomib resistance in multiple myeloma.

**Highlights:**

- SNUPR allows simultaneous profiling of PERK, IRE-1 and ATF6 activation with single-cell resolution.
- Inhibition of protein synthesis via PERK control the activation levels of the IRE-1/XBP1s and ATF6 pathway.
- IRE-1 activation and associated transcriptional signatures predict the outcome of patients with multiple myeloma treated with Bortezomib.
- IRE-1 activity, but not PERK or ATF6, is essential to acquire bortezomib resistance in multiple myeloma cell lines.

**Graphical abstract:** 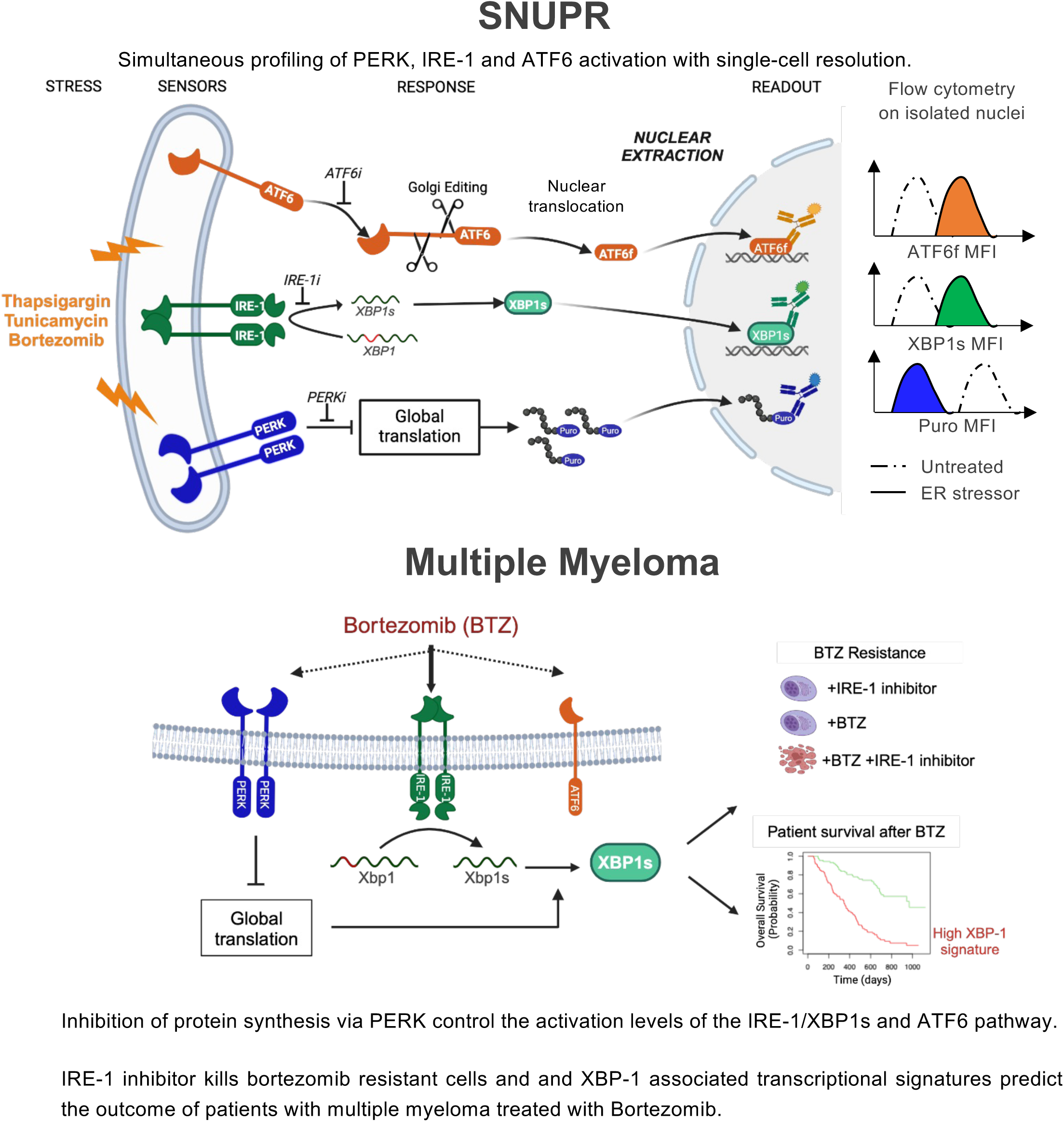

## INTRODUCTION

The tumour microenvironment contains biochemical stressors that can disrupt protein folding within the endoplasmic reticulum (ER) and affect cell viability. Accumulation of misfolded proteins in the ER triggers a cellular response known as the unfolded protein response (UPR)^1^. The UPR comprises distinct signalling branches, each activated by the dissociation in the ER lumen of the HSPA5 chaperone (BiP) from three ER-resident sensors. These sensors include activating transcription factor 6 (ATF6), inositol-requiring enzyme 1-alpha (IRE1α, ERN1), and (PKR)- like endoplasmic reticulum kinase (PERK, EIF2AK3). Each sensor triggers a unique cellular response and leads to the activation of specific transcription factors, ultimately modulating a range of genes involved in ER homeostasis^2^. (i) IRE1α splices the mRNA encoding for the transcription factor X-box binding protein-1 (XBP1) involved in the expression of genes regulating ER size and function. (ii) ATF6 exits the ER to reach the Golgi apparatus and undergoes proteolytic release in the cytosol as ATF6f, which in turn act as a transcription factor that induces the synthesis of *XBP1* and ER chaperone genes like *HSPA5* (BiP) and others. (iii) PERK activation mediates the phosphorylation of eukaryotic initiation factor 2-alpha (eIF2α). While eIF2α phosphorylation leads to a decrease of global protein synthesis, it also favors the translation of specific transcripts like activating transcription factor 4 (ATF4), C/EBP homologous protein (CHOP, DDIT3) or the phosphatase co-factor PPP1R15a (GADD34) mRNAs ^3,4^.

As a key regulator of proteostasis and cell death, the UPR has gained a lot of interest as a promising target in anti-tumoral therapy. Depending on the duration and degree of the ER stress, the UPR can provide either survival signals or trigger cell death through apoptosis. Tumour progression require high level of protein synthesis and exposes cells to multiple extrinsic and intrinsic stressors that can lead to chronic UPR activation and malignant progression^1,5^. On the other hand, non-cancerous cells can also show activation of UPR pathways but in general rely less on constant high levels of translation. This difference offers an advantage for potential chemotherapies, like proteasome inhibitors, by modulating the UPR to target cancer cells^6^. Sustained pharmacological induction or repression of the UPR could exert beneficial anti-tumoral effects. Henceforth the interest to combine standard therapies with drugs directed towards unresolved ER stress or UPR modulation to restrain tumour growth; some of which have already shown to be effective in pre-clinical tumour models^7–10^.

Our understanding of the interplay between UPR effectors, how it influences the balance between cell survival and cell death; and our capacity to predict ER-stress responses in physiological contexts remains limited due to technical limitations. We developed “Single Nuclei Analysis of the Unfolded Protein Response” (SNUPR), a method that allows the simultaneous measure of ATF6f, IRE-1/XBP1s and PERK pathways activation, as well as their interplay on a single-cell and time-resolved basis.

Using SNUPR, we were able to highlight the heterogeneity of UPR activation in response to standard ER stressors on different cancer cell lines. This allowed us to shed light on how the intensity and duration of translation inhibition via the PERK pathway can shape responses from the other UPR branches during ER-stress across various cancer cell lines. Additionally, our analysis allowed us to uncover the importance of the IRE1/XBP-1s axis in the resistance to bortezomib chemotherapy in multiple myeloma cell lines. Furthermore, transcriptomic analysis of multiple myeloma cohorts confirmed our hypothesis that the XBP1 gene signature alone can predict the outcome of multiple myeloma patients treated with bortezomib as monotherapy. By revealing the complex interplay and hierarchy between UPR branches via SNUPR, our findings underline how strategic manipulation of the UPR could present a promising therapeutic strategy for treating cancer or to stratify multiple myeloma patients to predict treatment efficacy.

## RESULTS

### Subhead 1: Correlation between ER-stress sensors activation and survival prognosis in acute myeloid leukaemia and breast cancer patients

Tumour growth often induces oxidative stress and glucose deprivation which in turn can cause oxidative damage and glycosylation defects that result in protein misfolding, ER stress and UPR activation^11^. The particular role of the different UPR branches in cancer is controversial, with both positive and negative outcomes described in the literature ^8,10,12–14^.

To gain insights whether UPR branches are associated with prognosis in human cancer, we stratified Acute Myeloid Leukaemia (AML) and breast cancer patients by their expression levels of ATF6f, XBP1s and ATF4-targeted genes (Figure 1A). We then evaluated patient survival over time generating Kaplan-Meier curves on those subgroups of patients defined on mRNA expression levels (Figure 1B, 1D and Supplemental Figure 1)^15^. Only a high expression of certain ATF6-associated genes was correlated with increased patient survival for both types of cancer, with 5 out of 14 target mRNAs for AML (Figure 1B and Supplemental 1A) and 3 out of 5 target mRNAs for breast cancer (Figure 1D) presenting significative difference. We also found that several target genes of XBP1s correlated with longer median survival in AML patients, 7 of which showing statistical significance (Figure 1D and Supplemental Figure 1B). Overall, patients with cells overexpressing genes that are transcriptionally-dependent on XBP1s and ATF6f showed cross-correlation of expression (Figure 1C and 1E), but not ATF4, and had a markedly enhanced survival rate.

**Figure 1.**
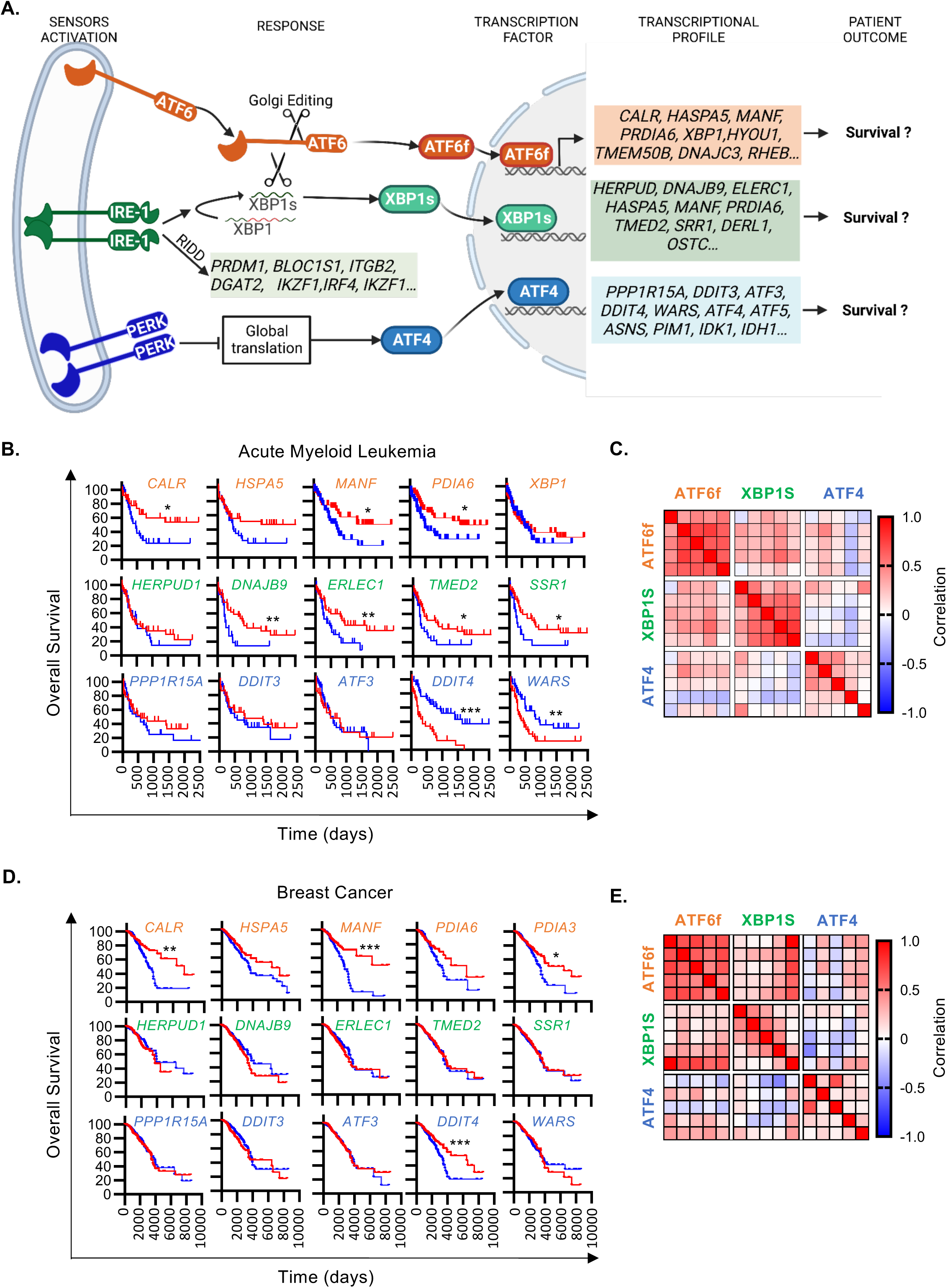
UPR signatures correlate with survival prognosis of myeloid leukemia and breast cancer patients. **(A)** Scheme of UPR three main branches. **(B-E)** Kaplan Meier survival curves and correlation matrices of genes under the control of ATF6, XBP1s or PERK corresponding to AML and breast cancer cohorts were generated using the Xena browser (USCC). Kaplan Meier curves were done based on dichotomized gene expression, specifically for values below quartile 1 (blue) and above quartile 3 (red) of both malignancies. Genes indicated are described to be under the control of ATF6 (orange), IRE-1/XBP1s (green) and PERK/ATF4 (blue). **(B)** Kaplan Meier curve and **(C)** correlation matrix for the AML cohort. **(D)** Kaplan Meier curve and **(E)** correlation matrix for the breast cancer cohort. Statistical analysis was performed using Log-rank test. \**P* < 0.05, \*\**P* < 0.01, and \*\*\**P* < 0.001.

Besides unconventional splicing of *XBP1* mRNA, another hallmark of IRE-1 activation is the cleavage of different RNAs through a process termed RIDD (Regulated IRE1- Dependent Decay), leading to degradation of mRNAs coding for genes such as *ITGB2* and *IKZF1* (Supplemental Figure 2) ^16–21^. In contrast to what we observed with XBP1 targets, we noted a variable and non-significant trend in survival rates when correlating survival with the expression level of RIDD targets (supplemental Figure 2) in AML. These results imply that the canonical IRE-1 activity, through *XBP1* splicing, is most likely contributing to patient survival in AML^16–21^.

Contrastingly, most of the PERK/ATF4 target genes, such as *PPP1R15A* (GADD34) and *DDIT3* (CHOP), showed no correlation or anti-correlation with survival prognosis (Figure 1B, 1D and Supplemental Figure 1). Nonetheless, certain non-exclusive ATF4 targets like *DDIT4, WARS*, *PIM*, and *PSAT1*^22^, were associated with lower survival rates in AML when overexpressed (Figure 1B and supplemental Figure 1C).

Altogether, the association of IRE-1/XBP1 and ATF6 signatures, but not ATF4 with survival suggests that in the presence of ER stress, UPR sensors may not be all activated simultaneously. Moreover, our result suggests that a cancer-specific association between IRE-1 and ATF6 activation and survival rates. Hence, a potential dichotomy and heterogeneity in ER stress sensors activation may influence disease progression and therapeutic response.

### Single cell resolution profiling of UPR signalling branches via SNUPR

The heterogeneity and hierarchical activation of IRE-1, ATF6, and PERK in response to ER stress is not well understood. Despite multiple studies linking ER stress sensor activation with cell death, the specific effects of proteasome inhibitors, such as bortezomib^23^, on this process remain poorly understood. Additionally, the use of transcriptional approaches to investigate this phenomenon can be misleading due to the central role of post-transcriptional regulation in ER stress responses. We developed SNUPR (Single Nuclei UPR profiling), a flow cytometry-based method that allows to delineate the activation of all three UPR branches, ATF6, IRE-1, and PERK with single-cell resolution. SNUPR uses multi-parametric flow cytometry of single nuclear suspensions to measure XBP1s and ATF6f nuclear translocation as well as intracellular puromycin incorporation^22,24,25^ to simultaneously assess transcription factor translocation with overall inhibition of mRNA translation as an indirect measure of PERK activation (Figure 2).

**Figure 2.**
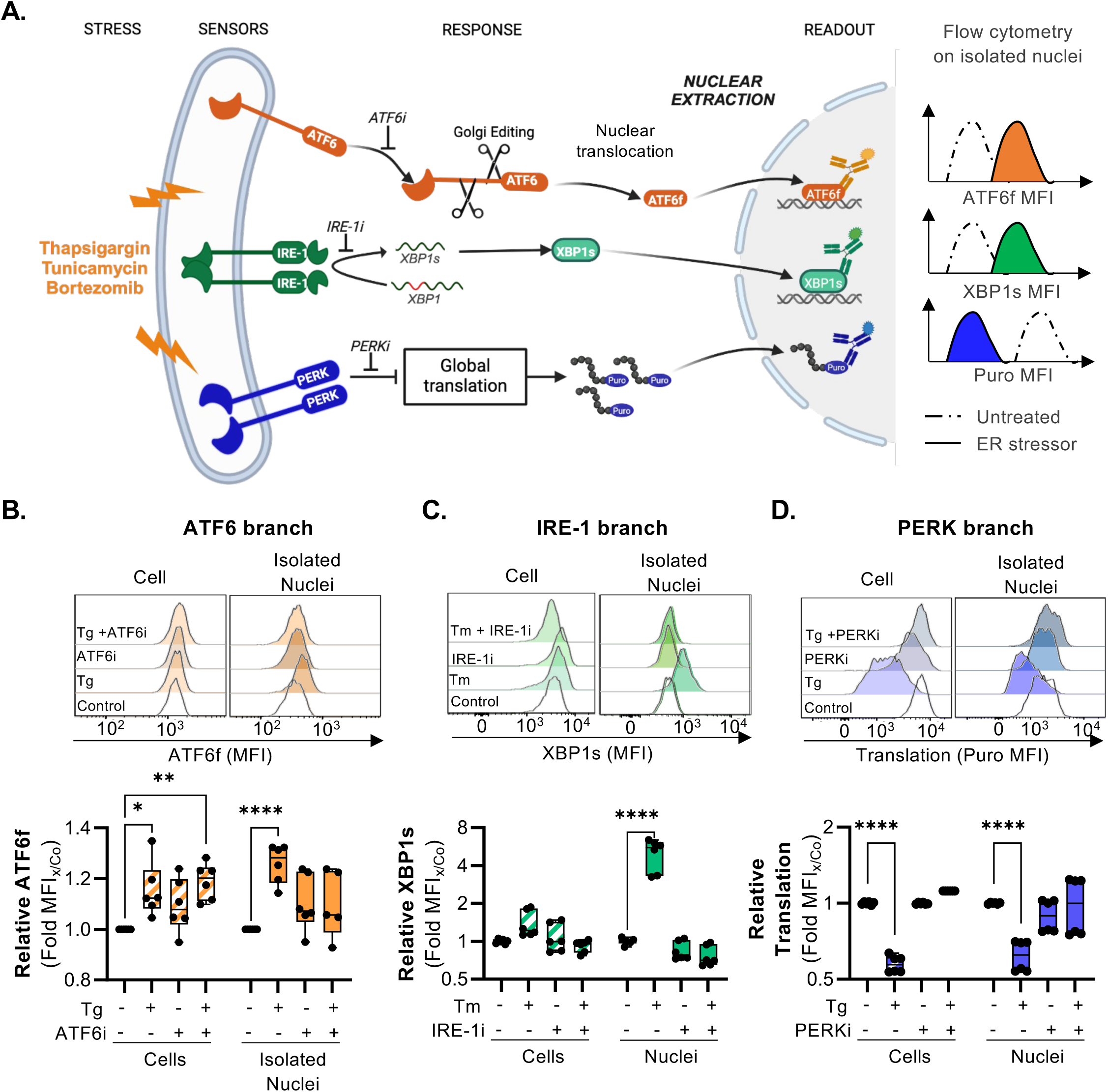
SNUPR, a method to profile sensors activation during ER stress. **(A)** Scheme of SNUPR method. Following nuclei extraction, the activation of UPR branches is profiled by flow cytometry measurements of XBP1s and ATF6f translocation as well as puromycin levels as a readout for protein synthesis. **(B)** THP1 were treated with 400nM of thapsigargin for 4h in presence (or absence) of the ATF6 inhibitor CeapinA7 (6µM). Afterwards, nuclei were extracted and ATF6f levels were analysed on both permeabilized nuclei and cells. **(C)** THP1 were treated with 100ng/mL of tunicamycin for 6h in presence (or absence) of the IRE-1 inhibitor 4µ8c (10µM). Afterwards, nuclei were extracted and XBP1s levels were analysed on both permeabilized nuclei and cells. **(D)** Hela cells were treated with 400nM of thapsigargin for 30min in the presence (or absence) of the PERK inhibitor GSK2656157 (100µM) and treated with puromycin for 15min. Afterwards, nuclei were extracted and puromycin levels were analysed on both permeabilized nuclei and cells. Statistical analysis was performed using Mann–Whitney test (\**P* < 0.05, \*\**P* < 0.01, \*\*\**P* < 0.001 and ****P ≤ 0.0001).

First, we isolated nuclei from plasmacytoid dendritic cell line (CAL-1) and monocytic AML cell line (THP-1) cells and used microscopy to confirm the enrichment of nuclei post-extraction, further validating their purity through immunoblot and flow cytometry assays targeting nuclear, plasma membrane and other organelles (supplemental Figure 3A- D). There was minimal presence of plasma membrane, mitochondrial, lysosomal, and endoplasmic reticulum contaminants in nuclear suspensions, indicating that our protocol allowed efficient nuclei isolation with minimal contamination from other intracellular compartments, permitting accurate quantification of transcription factors translocation.

Next, we employed fluorescent labelled antibodies to monitor the translocation of XBP1s and ATF6f transcription factors by flow cytometry (Figure 2A). We extracted nuclei from ER stress-induced THP-1 cells, treated with or without thapsigargin (Tg) or tunicamycin (Tm), in the presence or absence of specific inhibitors of IRE-1 (4µ8c)^26^, SP1/2 that mediate ATF6-cleavage (CeapinA7)^27^ or PERK (GSK2656157)^28^. By comparing the staining of the transcription factors between cells and nuclei, we obtained an enhanced signal to noise ratios for both XBP1s and ATF6f post-nuclear extraction (Figure 2B-C). The inhibition of UPR signalling pathways resulted in a decrease in nuclear staining of the corresponding transcription. These results demonstrate, on one hand the specificity of the measurements, and on the other, the higher signal to noise ratios obtained when analysing nuclear extractions; further corroborating the effectiveness of SNUPR in monitoring IRE-1 activation and ATF6 cleavage.

Our efforts to find suitable antibodies for detecting ATF4, as a direct readout of PERK activation by flow cytometry were however unsuccessful, and we turned towards protein synthesis inhibition measurement instead^29^ (Figure 2D). Puromycin is a tRNA-aminoacyl analogue and measuring intracellular puromycin incorporation into peptides, we monitored global translation inhibition mediated by PERK-dependent eIF2α phosphorylation^22,24^. We observed a very strong and significant correlation between nuclear and cellular levels of puromycinilated peptides^30^ (supplemental Figure 3E). A marked decrease in puromycin signal was observed in nuclei from cells treated with the ER stressors Tg and Harringtonine (Figure 2D and Supplementary Figure S3E). Upon PERK inhibition (GSK2656157, PERKi), the puromycin signal remained unaltered after Tg treatment, supporting the validity and specificity of our approach as readout for PERK activation (Figure 2D). Consequently, SNUPR, by monitoring XBP1s and ATF6f translocation together with puromycin incorporation on isolated nuclei, allows the simultaneous measure of all three UPR signalling branches activation with high accuracy.

### Deciphering cell type- and stressor-dependent UPR heterogeneity using SNUPR profiling

Given the clinical relevance of a potential association between UPR heterogeneity and patient response to treatment and survival, we treated seven human cell lines of diverse origin with thapsigargin (Tg) and tunicamycin (Tm) and applied SNUPR and qPCR analysis to monitor UPR activation over time (0-4h) (Figure 3 and supplemental Figure 4). SNUPR revealed divergent UPR activation patterns in response to the two stressors, but also great variations among the cell lines tested with the same stressor (Figure 3 and Supplemental Figure 4A). Solely based on the duration and intensity of the translation inhibition driven by the two compounds, we identified three distinct response patterns (Figure 3A). Group 1 (MOLT-4 and U937 cells) showed complete translation arrest within 30min to 4h of Tg treatment. Group 2 (HeLa, HEK293T, KASUMI and CAL-1) underwent up to 70% translation inhibition after 30min, recovering to initial levels within the subsequent 3h. Lastly, Group 3 (THP-1 monocytic AML line) presented little to no reduction of translation during Tg treatment, mirroring the previously reported behaviour of murine dendritic cells^31^.

**Figure 3.**
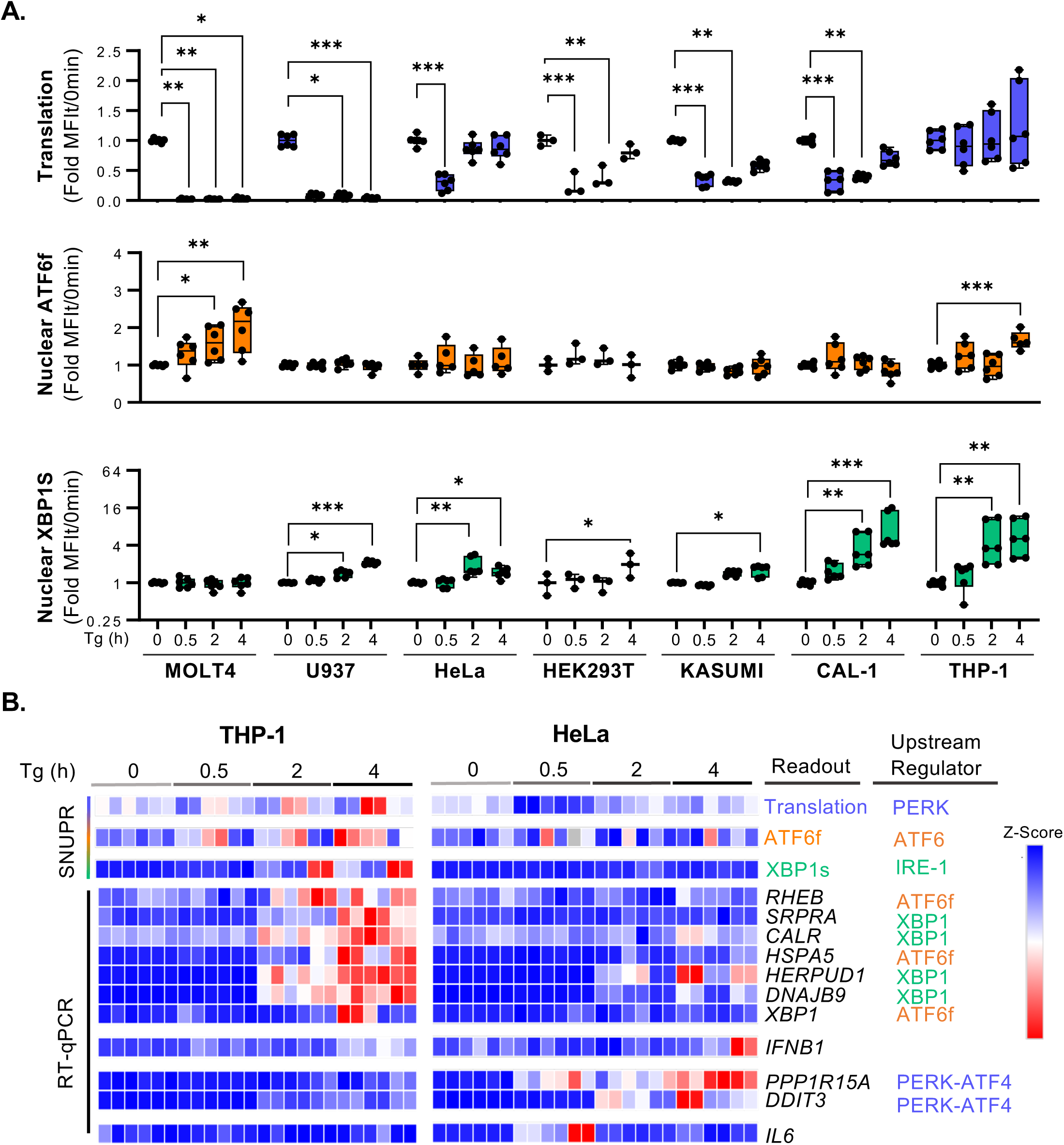
Induction of acute ER stress induces different UPR profiles in cell lines. **(A)** 7 different types of cancer cell lines were treated with thapsigargin (Tg) for 30min, 2h or 4h to induce acute ER stress prior to nuclei extraction and SNUPR profiling of UPR activation. Shown are boxplots of MFI fold changes (MFIt/t0 min) values representing translation level, ATF6f and XBP1s translocation. **(B)** Heatmap representation of SNUPR measurement as well as the expression level of ATF4-, ATF6- and XBP1s-target mRNAs measured by RT-qPCR. Each column corresponds to one duplicate of three independent experiments. Statistical analysis was performed using Kruskall-Wallis test for each cell line. \**P* < 0.05, \*\**P* < 0.01, and \*\*\**P* < 0.001.

Upon examining IRE-1 and ATF6 activation patterns, a significant increase in XBP1s translocation was noted within 4h of Tg treatment across most cell types, but a significant increase of ATF6f was only detected in two of the seven cell lines (MOLT- 4 and THP-1) (Figure 3A). Regarding translation, all cell lines except THP-1 cells displayed a significative reduction of protein synthesis after 30min, in some cases beginning to recover after 2h of Tg treatment (Figure 3A and 3B).

To determine if the different patterns of UPR were solely dependent on the cell line, we decided to use other ER-stress inducer. When Tunicamycin was used as the stressor, we observed a different pattern of response (supplemental Figure 4A). While most cells increased XBP1s expression after 4h, no significative changes were found on ATF6f, except for the U937 cell line. As for translation inhibition, all cell lines analysed displayed a moderate translation decrease within 4h of Tm treatment. Notably, THP-1 cells displayed increased translation within the first 30min of treatment, before returning to basal levels (supplemental Figure 4A). To validate SNUPR observations, we assessed the induction of mRNA levels of target genes of XBP1s, ATF6f and ATF4 in HeLa and THP-1 cells, since these cells displayed contrasting UPR induction patterns (Figure 3B and supplemental Figure 4B). The RT-qPCR results mirrored SNUPR results, with THP-1 cells experiencing a significant increase in ATF6f and XBP1s specific transcripts, such as *XBP1, HSPA5, RHEB,* or *CALR*, under Tg-induced stress; while HeLa cells primarily induced ATF4-dependent transcription of *DDIT3* (*CHOP*) and *PPP1R15A* (*GADD34*) (Figure 3B and supplemental Figure 4B).

We next wondered whether the level of expression of each ER stress sensors could reflect the pattern of activation of the different cell lines. However, levels of IRE-1 or PERK measured by immunoblot (supplemental Figure 5A) did not show any significant correlation with the capacity of the cells to block translation or translocate XBP-1 upon ER-stress (supplemental Figure 5B). This highlights the challenge of predicting functional cellular responses based solely on steady-state phenotypic markers, and supports the advantage of using functional readouts such as the ones measured in SNUPR to follow UPR activation dynamically and dissect the interplay among the three different individual UPR branches.

Interestingly, we observed that the absence and presence of *PPP1R15A* and *DDIT3,* in THP-1 cells and HeLa cells respectively, coincided with the degree of measurable translation arrest in response to the stressors (Figure 3B), where THP-1 cells did not block protein synthesis, HeLa cells did. Additionally, cells whose protein synthesis remain most active during stress induce more nuclear translocation of XBP1s, while the opposite trend is observed with ATF6f (supplemental Figure 5C). Moreover, despite the rapid induction of *XBP1* mRNA splicing 2 hours post-Tg treatment in both HeLa and THP-1 cells (supplemental Figure 4B), significant increases in nuclear XBP1s levels were only observed in THP-1 cells that did not block protein synthesis. Altogether, these results suggest that the magnitude of IRE-1/XBP1s response is inversely proportional to the degree of translation inhibition experienced by stressed cells (supplemental Figure 5D). Specifically, we hypothesized that the translation of *XBP1* and *ATF6* mRNAs might be hindered by PERK/P-eIF2α-mediated translation inhibition, thereby delaying XBP1s synthesis and downstream transcription of its target genes. In contrast, ATF6f appeared to be relatively less dependent on active translation (Supplemental Figure 5C), as its early mechanism of activation relies on the proteolytic cleavage of pre-existing ATF6 rather than its de novo synthesis. In conclusion, SNUPR enabled us to uncover a heterogeneous dynamic of UPR activation among distinct cell types and in response to different stressors, hinting a potential dependence of UPR responses on intensity and duration of PERK-p-eIF2α-mediated translation inhibition.

### Impact of translation arrest on activation of IRE1α/XBP1s and ATF6 Pathways

To further explore the effect of PERK-mediated translation arrest on nuclear XBP1s and ATF6f, we took advantage of the absence or presence of transient translation inhibition in THP-1 and HeLa cells, respectively. To test our hypothesis, we forced translation inhibition in THP-1 (Figure 4A) and blocked translation inhibition in HeLa cells upon ER-stress and measured the level of activation of the IRE- 1/XBP1 pathway. Although THP-1 and HeLa cells display contrasting UPR dynamics, both cell types express high levels of XBP1s after 4h of stimulation (Figure 3 and Figure 4). Co-treatment of THP-1 cells with Tg and the translation inhibitors rocaglamide (RocA) or cycloheximide (CHX) efficiently suppressed translation and reduced nuclear translocation of XBP1s and ATF6f after 4h of UPR (Figure 4A). As expected, HeLa cells underwent translation arrest very rapidly within 30min of Tg exposure (Figure 4B), although full recovery was observed within 4h. Pre-treatment of HeLa cells with the ISR inhibitor (ISRIB), a compound known to bypass the inhibitory effect of PERK-dependent eIF2α phosphorylation on translation^32^, circumvented this transient inhibition (Figure 4B) and led to significantly increased nuclear XBP1s levels at 4h post-treatment. ATF6f showed a similar trend of increased nuclear levels at 4h, but these were not statistically significant. To further investigate, the downstream effects of altering translation levels we measured XBP1s-dependent genes expression such as *RHEB*, *HERPUD1*, or *SRPRA* on Tg- and ISRIB-treated HeLa cells and detected no significant differences in mRNA expression levels (supplemental Figure 6A). These results suggest that the transient (30 min) inhibition of translation and delay in XBP1s protein expression observed in HeLa cells, can be circumvented by ISRIB, but is not prolonged enough to strongly impact the transcription of XBP1s target genes.

**Figure 4.**
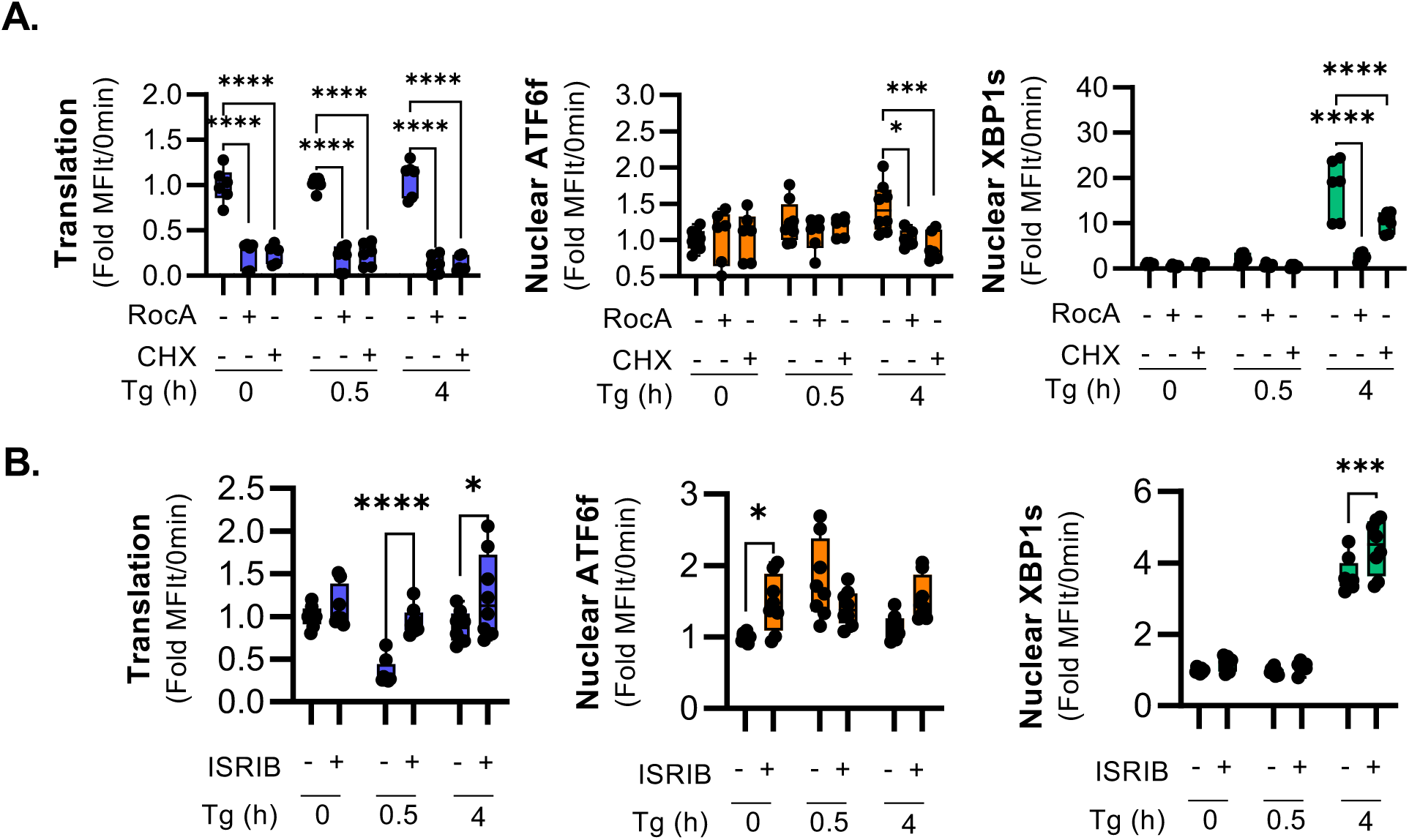
Translation arrest delays the activation of the IRE1/XBP1s and ATF6 pathways. THP-1, HeLa cells and peripheral blood mononuclear cells (PBMC) from healthy donors were treated 30min or 4h with thapsigargin (Tg, 400nM) in the presence or absence of different translation inhibitors prior to SNUPR profiling by flow cytometry. **(A)** Translation levels and translocation of ATF6f and XBP1s measured on THP-1 cells after treatment with Tg in combination with Rocaglamide (RocA, 100nM) or cycloheximide (CHX, 5uM). **(B)** Translation levels and translocation of ATF6f and XBP1s measured on HeLa cells after treatment with Tg combined with ISRIB (1ug/mL). Statistical analysis was performed using 2way ANNOVA test. \**P* < 0.05, \*\**P* < 0.01, and \*\*\**P* < 0.001.

### SNUPR highlights a cell-type dependent UPR profiles in PBMCs

We extended our observations of UPR activation to primary immune cells. For this, we expanded the capabilities of SNUPR by incorporating intracellular staining of lineage-specific transcription factors such as PU.1 (Monocytes) and GATA3 or BCL-6 (T cells) as well as global levels of epigenetic marks (supplemental Figure 7A-B). This enabled the dissection of UPR activation in heterogeneous primary cell samples such as human PBMCs. To validate this approach, we first stained healthy donor PBMC (whole cells) using surface and nuclear markers simultaneously (supplemental Figure 7A-B). Among the different nuclear markers examined, we found that PU.1 and GATA3 were sufficient to unequivocally identify monocytes, B cells and a third cluster of T/NK cells (supplemental Figure 7C). Using this strategy, we analysed isolated nuclei from PBMC samples treated with Tg in the presence or absence of guanabenz (GBZ), a molecule that inhibits eIF2α de-phosphorylation and thus inhibits the recovery of translation^33^ (supplemental Figure 7D). Our findings indicate that UPR sensors activate differentially in distinct cellular subsets in response to Tg. In addition, ATF6f was primarily observed in nuclei from monocytes and B cells, while XBP1s was predominantly found in monocyte and to a lower extent in B and T/NK cell nuclei (supplemental Figure 7D). Furthermore, and concordant with our previous results on cell lines, primary monocytes displayed increased XBP1s levels after 4h of Tg activation, which was significantly reduced after translation inhibition with GBZ (supplemental Figure 7D).

Taken together, these findings validate that translation arrest due to PERK activity can modulate IRE-1/XBP1s axis responses. We further show that SNUPR, in combination with specific nuclear lineage markers, offers an effective method to profile the activation of the three UPR branches in mixed primary cell samples.

### Role of IRE-1/XBP1s signalling in Bortezomib resistance in multiple myeloma cells

Upon chemotherapies resistant cancer cells can induce or rely specific transcriptional programs^34^. We sought to further explore how heterogeneity in UPR activation might impact the chemotherapy response in patients with multiple myeloma. Specifically, we focused on the role of endoplasmic reticulum (ER) stress response in mediating the efficacy of Bortezomib (BTZ) (Velcade, previously PS- 341). BTZ is a proteasome inhibitor that is currently included into the first line of treatment against multiple myeloma (MM) and has been reported to induce cytotoxic ER stress^35^.

We used SNUPR to profile the UPR response to BTZ treatment across several leukemia and MM cell lines over time (0-6h). BTZ treatment resulted in decreased protein synthesis in most cell lines, with the notable exception of U937 cells (supplemental Figure 8A). This contrasted with the results observed upon Tg treatment (Figure 3). Although variable in intensity, XBP1s translocation was consistent throughout cell lines; ATF6 translocation, however, was only detected on the leukemia cell lines but not in RPMI and LPI MM cell lines (supplemental Figure 8A). Additionally, we used SNUPR to analyze nuclear accumulation of CHOP (DDIT3), a pro-apoptotic transcription factor commonly associated with the UPR and PERK-ATF4 pathway. CHOP induction by BTZ was induced in most cell lines, except for THP1, where only a moderate induction trend without statistical significance was noted. Taken together, these observations corroborate a BTZ-mediated activation of the UPR.

To assess whether activation of any of the specific UPR branches played an essential role in BTZ toxicity, we quantified BTZ-induced cytotoxicity after 24h of treatment in presence of specific pharmacological inhibitors for each of the three ER stress sensors. Inhibition of PERK, IRE-1 and ATF6 pathways did not rescue BTZ-induced cell death (Figure 5A, Supplemental Figure 8B). Moreover, in LP1 MM cells, a BTZ resistant subpopulation persisted even at higher doses of BTZ (Figure 5A, supplemental 8B and 8C, bottom). Interestingly, IRE-1 inhibition in this resistant cell subset significantly increased cell death, revealing that its survival depends on IRE-1 (Figure 5A; and supplemental 8B and 8C). Inhibition of IRE-1 with the RNAse inhibitor 4µ8c alone did not show any toxicity in absence of BTZ, while higher doses of the IRE-1 kinase inhibitor Kira6 decreased the overall survival of MM cells in all conditions (Figure 5B). Given the differences of specificity of the 2 compounds and knowing that Kira6 also inhibits the p38 and ERK MAP kinase ^36,37^, we suspect an IRE1- independent effect in Kira6 cytotoxicity at these higher concentrations. Overall, these results suggest that activating the UPR is not essential for BTZ to mediate cytotoxicity, but in contrast, in Multiple Myeloma cells, IRE-1 activity contributes to resistance to Bortezomib.

**Figure 5.**
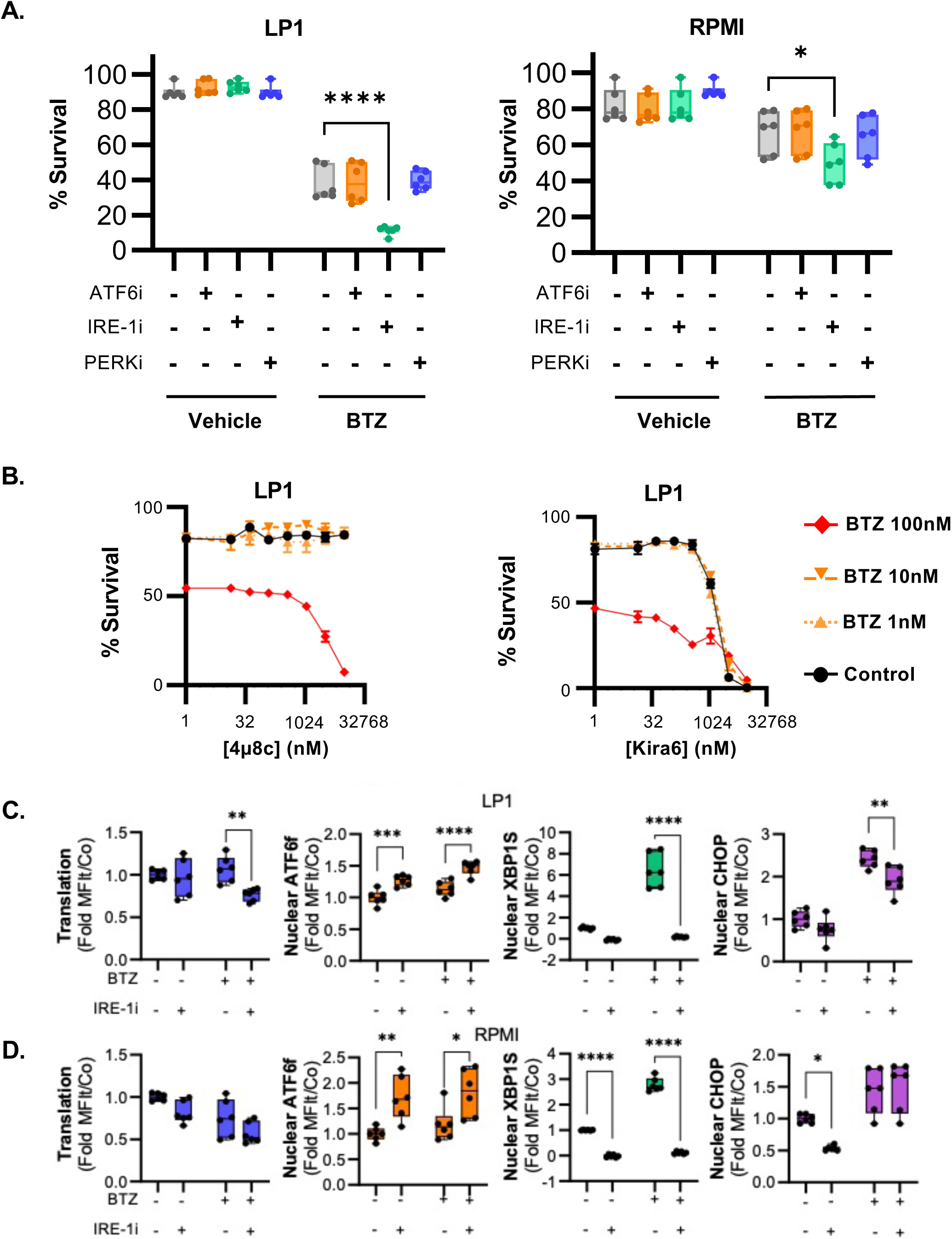
IRE-1 contributes to bortezomib resistance in multiple myeloma cell lines. Cell survival analysis were performed on MM cells to assess potential synergic effects between BTZ and UPR inhibitors **(A)** Flow cytometry analysis of survival of LP1 and RPMI MM cell lines after treatment with BTZ in combination with the ATF6 inhibitor (Ceapin A7, 6uM), IRE-1 inhibitor (4u8c, 10uM) or PERK inhibitor (GSK2656157, 100nM) for 24hr. **(B)** Cell survival analysis by flow cytometry of LP1 cells treated for 24h with different concentrations of BTZ (1nM, 10nM, 100nM) in presence of different concentrations of IRE-1 inhibitors 4µ8c (left) and Kira6 (right). **(C-D)** SNUPR profiling of **(C**) LP1 and **(D)** RPMI cell lines treated with BTZ 100nM for 4h in the presence or absence of IRE-1 inhibitor 4u8c (10uM). Statistical analysis was performed using 2- way ANOVA test. \**P* < 0.05, \*\**P* < 0.01, and \*\*\**P* < 0.001.

To further characterize the UPR mechanism of resistance to BTZ, we performed SNUPR and transcript analysis on LP1 and RPMI cell lines treated with BTZ in the presence or absence of 4µ8C (IRE-1i) (Figure 5C and 5D and supplemental Figure 9). Our results confirmed that IRE-1 inhibition decreases basal level of nuclear XBP1s, together with *XBP1* mRNA splicing and expression of XBP1s-regulated genes such as *HERPUD1* or *DNAJB9* (Supplemental Figure 9A). In contrast, ATF6f translocation was increased after IRE1 inhibition, but transcription of *ATF6a* mRNA and of its target *HSPA5* remained stable across all conditions (Figure 5C and 5D and Supplemental Figure 9A), suggesting that the modest effect of IRE1i on ATF6f translocation has only limited effect on downstream ATF6- dependent transcription and confirming that IRE-1/XBP1s is the only branch of the UPR that contributes to increase resistance to BTZ of MM cells.

### Pharmacological UPR modulation modestly contributes to overturn resistance to BTZ in multiple myeloma cell lines

Upon our findings of the significant impact of translation inhibition on the IRE-1/XBP1s axis and the involvement of this pathway in resisting to BTZ treatment, we harnessed these results by investigating whether UPR inducing drugs could synergize with BTZ. A reinforced UPR could further drive PERK activation and eIF2α phosphorylation levels to reduce XBP1s translation and thus undermine BTZ resistance by dwarfing the consequences of IRE-1 signaling (Figure 4).

We tested HA15, a specific inhibitor of the ATP dependent chaperone BiP/HSPA5, which was reported to exert its activity by inducing a lethal ER stress, particularly in melanoma cells^38,39^. We performed kinetic SNUPR on LP1 and RPMI cell lines treated with HA-15 over 8h (Figure 6A). HA15 triggered XBP1s translocation in both cell lines, as well as, some expected CHOP synthesis and translocation^39^. Despite CHOP activation, HA15 had no impact on translation levels in LP1 and even slightly elevated them after 8h in RPMI cells. We nevertheless tested the toxicity of HA-15 in MM cells, in presence of the different UPR inhibitors (Figure 6B). We found that HA15 efficiently killed both cell lines at concentrations above 5µM, but surprisingly inhibition of the IRE-1 pathway, and not of the other UPR branches, had a protective effect and reduced the efficacity of the HSPA5 inhibiting compound. Thus, contrasting with BTZ, HA15 seems to rely on IRE-1 activation to kill MM cells and inhibition of this specific UPR branch leads to enhance survival. These contrasting results called for further assessment of the combinatorial effect of exposing MM cells to HA15 together with BTZ. Co-treatment with both compounds had a modest enhancing effect on the activation of the UPR monitored with SNUPR (Figure 6C). Levels of translation were slightly reduced and accompanied with an elevation of CHOP nuclear levels and equivalent translocation of ATF6f and XBP1s (Figure 6C). When examining cytotoxicity, no synergy in the killing of MM cells could be observed by gradually increasing drugs quantities (Figure 6D), while HA15 killed efficiently BTZ resistant MM cells at micromolar concentrations. Interestingly, although the two drugs induced IRE-1 activation, it had opposite consequences for the cells: IRE-1 activity mediating survival in BTZ, while promoting cell death in response to HA15. Taken together, these findings suggest that the molecular mechanisms by BIP inhibition and Proteasome inhibitors induce cytotoxicity is different in MM and involves opposite roles of IRE-1 (Figure 5 and Figure 6).

**Figure 6.**
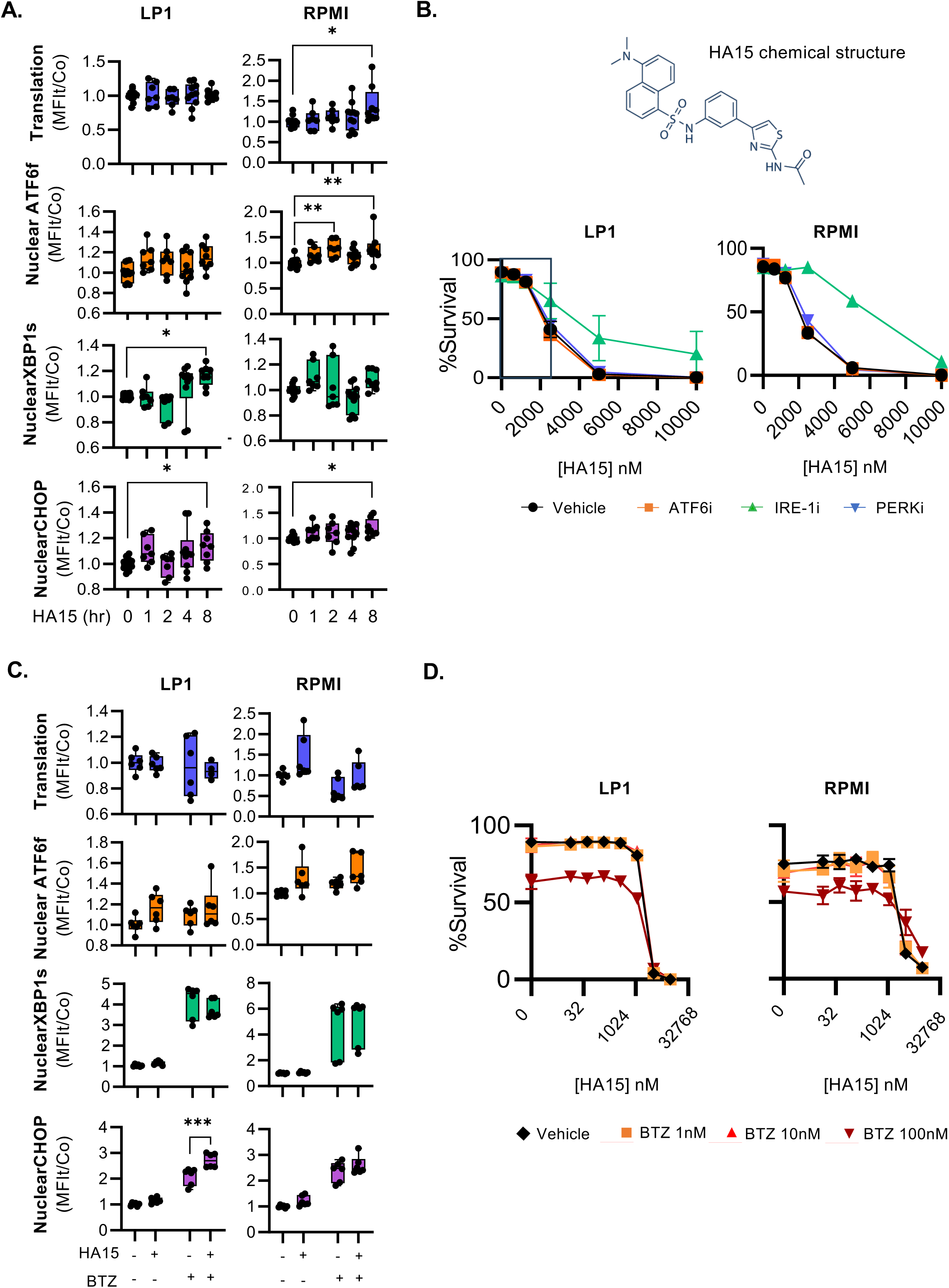
**BIP inhibition by HA15 induce cell death in BTZ-resistant and non-resistant multiple myeloma lines**. **(A)** Kinetic SNUPR of LP1 and RPMI cells treated or not with HA15 (5µM). **(B)** Flow cytometry analysis of survival of LP1 and RPMI cells treated with or without ATF6, IRE-1 or PERK inhibitors in combination with increasing concentrations of HA15 for 24h (0, 625nM, 1250nM, 2500, 5000 and 10000nM). **(C)** SNUPR profiling of LP1 and RPMI cells treated 4h with BTZ in presence or absence of HA15. **(D)** Flow cytometry analysis of survival of LP1 and RPMI cells treated with different concentrations of BTZ (1nM, 10nM and 100nM) in combination with increasing concentrations of HA15 for 24h.

### UPR Gene Signatures and Their Clinical Relevance in Multiple Myeloma

We have shown that the IRE-1/XBP1 pathway is associated with resistance to BTZ treatment in human MM cell lines in-vitro. To test the clinical relevance of these findings, we investigated whether the presence of XBP1 transcriptional signatures correlated with resistance to BTZ and/or survival in treated MM patient. For this analysis, we focused on the Mulligan cohort of relapsed MM patients who were treated exclusively with BTZ^40^. Patients were stratified based on the expression levels of gene signature consisting in a set of XBP1 target genes prior to treatment, and survival probability was monitored over time (Figure 7). Notably, high mRNA expression levels of most genes of the IRE-1α/XBP1s pathway significantly correlated with a poor prognosis (Figure 7A). To determine whether the XBP1 gene signature is specifically predictive of poor prognosis following BTZ treatment, or whether it serves as a marker of adverse outcomes independently of the treatment, we conducted an analysis using three different MM patient cohorts of patients treated with Dexamethasone, with anti-CD38 as monotherapy, or receiving a combination of immunomodulatory drugs (IMiDs), dexamethasone (Dex), and high-dose melphalan followed by autologous stem cell transplantation (Figure 7B, 7C and 7D; respectively). Strikingly, the XBP1 gene signature had strong predictive value in patients treated with Bortezomib, but not in patients treated with other treatments. This association reinforces the potential relevance of the IRE-1/XBP1s pathway in BTZ resistance and the potential therapeutic value of its modulation (Figure 7E).

**Figure 7.**
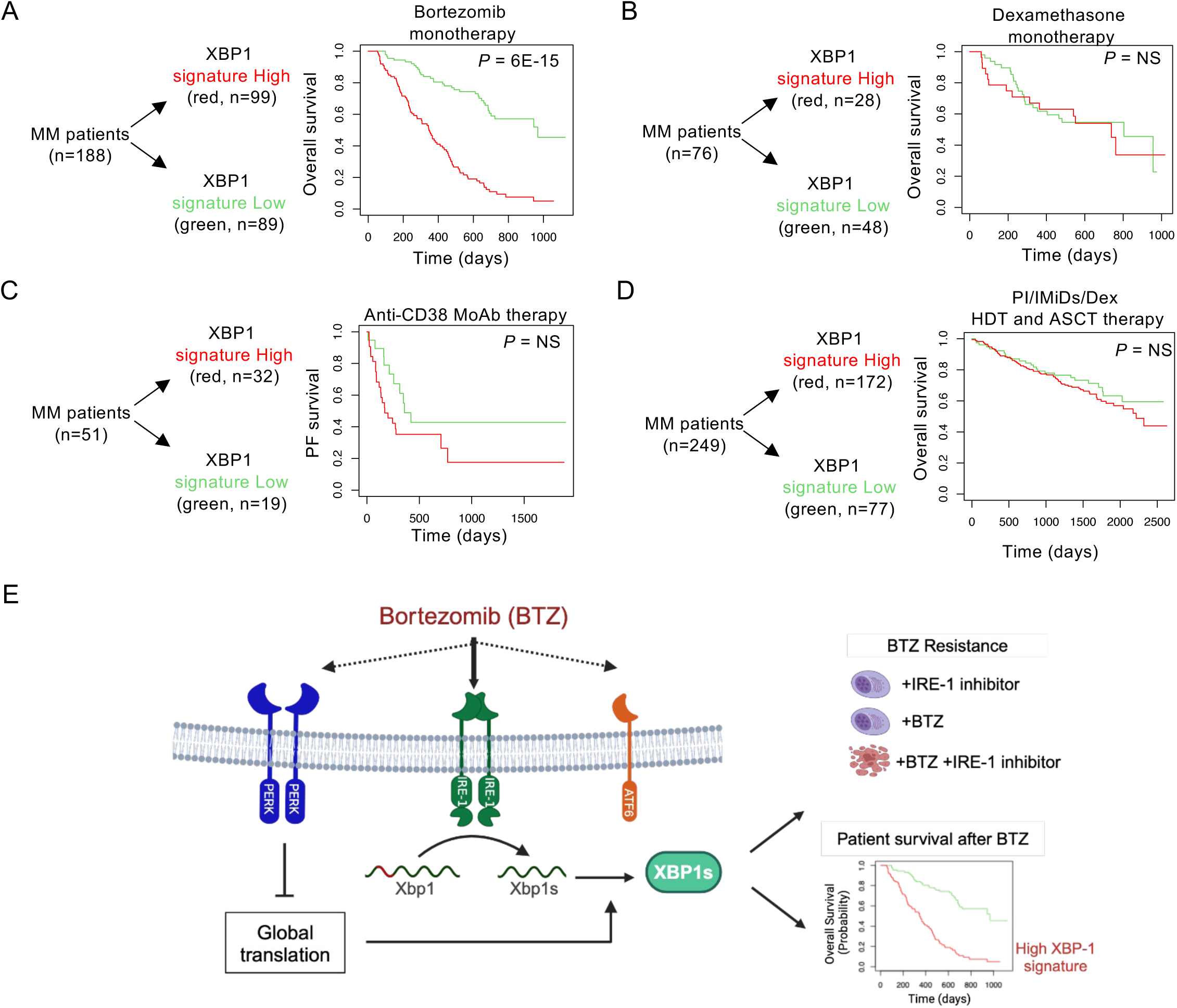
XBP1 gene signature correlates with unfavorable outcomes in Multiple Myeloma patients treated with Bortezomib. Transcriptional levels of XBP1 target genes was used to stratify and perform survival curves of cohorts of MM patients treated with Bortezomib (A), Dexamethasone (B), anti-CD38 monoclonal antibody (C) and PI/IMiDs/Dex HDT and ASCT therapy (D). Summary of results and model (E)

## DISCUSSION

Under physiological and pathological conditions, cells can activate the unfolded protein response (UPR) to cope with the accumulation of unfolded or misfolded proteins in the ER. The primary purpose of the UPR is to alleviate cellular stress by employing various biochemical mechanisms that enhance ER protein folding capacity, degrade misfolded proteins and suppress general protein synthesis to aid cell survival; or activate cell death programs if there is sustained stress^2^. Besides rapid proliferation and high mutation rate, tumour cells are often confronted with hostile environmental conditions, such as hypoxia, nutrient depletion, and presence of free radicals, which makes them highly reliant on the UPR to thrive^5,41^ and resist to chemotherapy. This makes of the UPR an interesting target pathway for therapeutic approaches aimed at eliminating cancer cells.

Previous methods lacked the ability to simultaneously measure individual UPR branch activations, limiting our understanding of UPR dynamics and inter-pathway crosstalk. We developed SNUPR, which innovatively combines nuclear extraction, with flow cytometry and translation measurements to quantify branch-specific UPR activation at a single-nuclei resolution, significantly enhancing the analysis of UPR pathways^2^, like XBP1s, ATF6f and CHOP after nuclear translocation, along with active translation through measurement of puromycin incorporation into cells^24,30^. Decreased puromycinylation represent a direct readout of PERK-dependent inhibition of protein synthesis. By purifying nuclei rather than permeabilized cells, we increased resolution and contrast to simultaneously monitor all these elements in different types of samples. Furthermore, by incorporating lineage-associated transcription factors, we show the potential of SNUPR to explore UPR activation in different cell subsets within heterogeneous cell populations such as PBMC. Further studies focusing on developing nuclear panels for spectral flow cytometry will allow to extract cell-type and state specific SNUPR profiles. Importantly, the resolutive power of SNUPR can be further enhanced or adapted for different biological models by utilizing or developing suitable monoclonal antibodies raised against cell-specific transcription factors or other UPR-dependent transcription factors such as ATF4.

SNUPR uncovered the heterogeneity of UPR responses on different cell types exposed to ER stressors, revealing differences between closely related cell types. Thapsigargin and tunicamycin are two classically used chemical ER stressors, both leading to misfolded protein accumulation in the ER lumen but through different mechanisms of action. Irrespective of the cell type analysed, SNUPR confirmed that Tg and elicit distinct ER stress responses and kinetic of action on different cell types, something of significance since these chemicals are generally used indistinctly for UPR studies.

Our observations also highlighted a hierarchy between the IRE-1/XBP1s and PERK pathways activation. This seem to be due to the translational control of XBP1s production by protein synthesis inhibition mediated by PERK. In this way, PERK controls the dynamics of UPR at the translational level independently of efficient IRE1-dependent mRNA splicing. Inhibitory eIF2α phosphorylation by PERK is counteracted by the UPR-inducible GADD34/PP1c phosphatase complex, which sets the pace for rescuing protein synthesis^22^. The balance between PERK and GADD34 can indirectly determine the levels of XBP-1 and ATF6 activation displayed by stressed cells. Notably, GADD34 has been shown to play a critical role in translation recovery within 4h after UPR activation^42,43^, coinciding with the emergence of XBP1s expression. Given the availability of numerous pharmacological inhibitors targeting the integrated stress response (ISR) pathway^44^, interference with PERK, eIF2α or GADD34 could be used to regulate not only the ISR, but also to modulate XBP1s signalling for therapeutic purposes. While BTZ effectively induces apoptosis in multiple myeloma, response to BTZ in patients is heterogenous and resistance often develops. Using SNUPR, transcriptomics and patient survival data, we identified a patient subset lacking XBP-1 signature showing enhanced response to BTZ. Further prospective clinical studies will be required to prove if our gene signature can be used as a personalized medicine tool to identify patients that can benefit of BTZ either as monotherapy or combined with lower doses of other treatments.

Bortezomib (BTZ) has proven effective in inducing apoptosis in multiple myeloma cells, significantly improving patient outcomes through modulation of several pathways, including endoplasmic reticulum stress signalling^45–47^. However, the persistent issues of frequent relapses and treatment resistance continue to limit its therapeutic success. It has been proposed that resistance may arise due to mutations in the highly conserved binding pocket within the proteasome subunit β5 (PSMB5)^48^. However, this explanation appears insufficient since mutations in the PSMB5 gene do not consistently account for BTZ resistance in primary MM sequencing studies^49,50^.

Current standard of care treatment of MM includes BTZ, but also Thalomide, Dexamethasone, high dose radiation therapy and autologous stem cell transplantation. One of the goals of personalized medicine is to avoid treatment of patients that will not benefit of a therapy, thus avoiding secondary effects. In our study, by combining stratification using the XBP-1 signature in patients treated with BTZ as monotherapy, we identified patients showing high survival rates, similar to the survival curves observed in patients treated with the standard of care combo therapy (Figure 7A vs 7D). Further studies and clinical trials will need to be performed to probe the use of SNUPR and XBP1s-dependent gene signature can identify patients that can benefit of BTZ as monotherapy, or in combination with lower doses of the combo therapy. Moreover, we observed that the combination of BTZ with IRE-1 inhibitor (4µ8C) eliminated proteasome inhibition resistant cells, thus suggesting that this XBP1 signature could be used to identify patients that would strongly benefit of treatment with IRE-1 inhibitors.

The UPR pathway has emerged as a key factor in BTZ resistance in MM. However, its exact role and the implication of its different branches in BTZ cytotoxicity and resistance have remained elusive^51^. Existing evidence presents contradictory contributions of the IRE-1 and PERK branches to BTZ resistance, underlining the complex and heterogenous role of the UPR in modulating cell survival and death in MM^52,53^. We hypothesized that this unresolved complexity could be attributed to the variability of the interplay existing among the different UPR signalling pathways, so we dissected the impact of BTZ on UPR dynamics in different cancer cell lines. BTZ treatment activated all three UPR branches in most leukaemia cells, except for MM cell lines.

Interestingly, MM cells failed to induce an ATF6f response, even though early ATF6 activation is relatively unaffected by PERK-dependent translation arrest. Although BTZ induced a robust UPR response and even CHOP translocation in MM cells, we observed resistance to cell death with escalating doses of the drug. Retrospective analysis of transcription profiles from BTZ-treated patients indicated that the presence of an IRE-1/XBP1s transcriptional signature, rather than a PERK/ATF4 signature, is strongly associated with poor clinical outcomes. The synergy observed between IRE1 inhibition and BTZ treatment in killing resistant MM cells further corroborated the relevance of this stratification analysis. Overactivation of the IRE-1 pathway appears therefore to contribute to BTZ resistance, contrary to previous reports suggesting that loss of XBP1s function or induction of the PERK pathway are major contributors to BTZ resistance^52–54^. The synergy observed between IRE1 inhibition and BTZ treatment in killing resistant MM cells further corroborated the relevance of this stratification analysis. Overactivation of the IRE-1 pathway appears therefore to contribute to BTZ resistance, contrary to previous reports suggesting that loss of XBP1s function or induction of the PERK pathway are major contributors to BTZ resistance^52–54^.

We compared the SNUPR profiling of BTZ with that of HA15, a specific inhibitor of BiP/HSPA5 shown to activate the UPR and kill effectively melanoma cells^38,39^. We observed that HA15 did not induce the UPR as strongly as BTZ, however, it still induced death in MM cell lines. Killing seemed to involve IRE-1 activity, rather than the PERK/ATF4/CHOP pathway and subsequent autophagy and apoptosis triggering, as suggested in initial reports cells^38,39^. The cytotoxic consequences of HA15 exposure may therefore vary considerably according to tumor cells specificity and the level of UPR heterogeneity displayed.

The IRE-1/XBP1s pathway appears as an important player on MM cell response to chemotherapeutic drug treatments and modulating its activity could offer new therapeutic strategies. The type of strategy chosen, however, will depend on the effect of the combined drug used over UPR pathways interplay, since we observed that the intensity of UPR activation, including the levels of translation inhibition, plays a significant role in the cytotoxic effects of ER stressors. BTZ and HA-15 for example, induce different UPR activation dynamics, and their cytotoxicity depends on opposite sides of the IRE-1 axis. These cell specific differences in the interplay between individual UPR signalling branches, and their potential contribution to chemotherapy, could only be evidenced with the resolutive capacity of SNUPR. This enhanced dissection of the UPR and its relationship to anti-tumoral drug treatment or resistance acquisition should prove useful in devising more effective MM treatments and predicting patient response.

In conclusion, our research offers new insights into the UPR’s role in haematological malignancies and its implications for overcoming chemotherapy resistance. By advancing the SNUPR technology and gene signatures, we pave the way for more targeted and effective treatments, underscoring the importance of dynamic and comprehensive pathway analysis in predicting treatment outcomes.

## MATERIAL AND METHODS

### Nuclear extraction

Prior nuclei extraction, the cells were stained with aqua dead from invitrogen for 20min on ice. Nuclei were extracted using EZ Prep (Sigma-Aldrich N3408) according to the commercial indication. Briefly, cells were harvested in 15mL tubes, washed with ice-cold PBS prior nuclei extraction and were centrifugated at 400g for 5 min at 4°C. Supernatant were removed and the cells were resuspended with 1.5mL of EZ. Nuclear suspensions were incubated on ice for 5 min, pelleted at 500g for 5 min at 4°C. Nuclei were resuspended in 1.5mL EZ lysis buffer, incubated on ice for 5 min, and pelleted before resuspension in 200µL of PBS 2% PFA for 20min on ice. Nuclei were pelleted at 500g for 5 min at 4°C and resuspend in the storage solution and conserved at -80°C until permeabilisation steps.

### Intracellular and nuclei staining for flow cytometry

Cells or nuclei were washed in cold PBS then fixed and permeabilized using FOXP3 fixation and permeabilization buffer (Thermofisher eBioscience) following manufacturer instructions. Cell and nuclei were blocked 10min at 4°C in blockage solution (Permeabilisation buffer 2%FCS) and stained with the following antibodies for 1h at 4°C in the dark (table 1). After incubation Nuclei were washed in FACS buffer prior flow cytometry analysis, on Canto II, LSR II UV BD cytometers and 5L Cytek Aurora spectral cytometer. The list of antibodies used can be found on Table 1.

**Table 1.**

| REAGENT or RESOURCE | SOURCE | IDENTIFIER |
| --- | --- | --- |
| Antibodies |  |  |
| Anti-human ATF6 A350(2358C) | R&D Biosystems | Cat#IC71527U |
| Anti-human XBP-1S PE(Clonc Q3-695) | BD bioscience | Cat#15802109 |
| Anti-puromycilated peptide A647(Clonc R4743L-E8) |  | RRID :<br>AB_2827926 |
| Anti-human CHOP A488 (B-3) | Santa Cruz | Cat#sc-7351 |
| Anti-human anti-nuclear pore (MAb414) | Biologend | Cat#682203 |
| Anti-human Calnexin (AF18) | Thermo scientific | Cat#MA3027 |
| ProLong™ Gold Antifade Mountant with DNA Stain DAP | invitrogen | Cat# P36931 |
| Anti-human SDHA | Abcam | Cat#ab14715 |
| Anti-human PERK | Cell signaling | Cat# 3192S |
| Anti-human GRASP55 | Proteintech | Cat#10598-1-AP |
| Anti-human βActin (AC-15) | MERCK | Cat# A5441 |
| Anti-human EIF2Bα | Invitrogen | Cat# PA5-28992 |
| Anti-human HDAC6 (H-300) | Santa Cruz | Cat#SC-11420 |
| Anti-human HDAC1 (10E2) | Cell signaling | Cat#5356S |
| Anti-mouse CD11b | BD Bioscience | Cat#562287 |
| Anti-mouse CD3 BV421 (Clonc: 145-2C11) | Biologend | Cat#BLE100335 |
| Anti-mouse Ly6C BV711 (Clonc: HK1.4) | Biologend | Cat#128037 |
| Anti-mouse NK1.1 BV510 (Clonc PK136) | Biologend | Cat#108373 |
| Anti-mouse CD4 APC-eFluor 780 (Clonc: RM4-5) | eBioscience | Cat#47-0042-82 |
| Anti-mouse MHC II BUV805 (Clonc M5/114.15.2) | BD Bioscience | Cat#748844 |
| Anti-human IRE-1 | Cell signaling | Cat# 3294S |
| Anti-human ATF6 | R&D Biosystems | Cat#IC71527U |
| LIVE/DEAD™ Fixable Aqua Dead Cell | Invitrogene | Cat#L34957 |
| Biological Samples |  |  |
| Murine blood C57BL/6J | The Jackson Laboratory | Stock 000664 |
| Chemicals, Peptides, and Recombinant Proteins |  |  |
| B-mercaptoethanol | VWR | Cat#0482-100ML |
| Cycloheximide (CHX) | Merck | Cat#01810-1G |
| Dimethyl sulfoxide (DMSO) | Merck | Cat#D8418-100ML |
| Harringtonine | Abcam | Cat#ab141941 |
| 4μ8c | Sigma-Aldrich | Cat#SML0949 |
| Ceapin A7 | Sigma-Aldrich | Cat#SML2330 |
| GSK2656157 | Sigma-Aldrich | Cat#5046510001 |
| Bortezomib | Sigma-Aldrich | Cat#5043140001 |
| Rocaglamide | Sigma-Aldrich | Cat#SML0656 |
| Thapsigargin | Sigma-Aldrich | Cat#T7458 |
| Tunicamycin | Sigma-Aldrich | Cat#SML1287 |
| Cycloheximide | Sigma-Aldrich | Cat#108-91-8 |
| Puromycin | Merck | Cat#P7255 |
| Critical Commercial Assays |  |  |
| Brilliant Stain Buffer Plus | BD Bioscience | Cat#566385 |
| Fixation/Permeabilization Solution Kit | BD Bioscience | Cat#554714 |
| Foxp3 / Transcription Factor Staining Buffer Set | eBioscience | Cat#00-5323-00 |
| LIVE/DEAD™ Fixable Aqua Dead Cell Stain (BV510) | Thermo Fisher | Cat#L34957 |
| Zombie UV™ Fixable Viability kit | Biolegend | Cat#423107 |
| Experimental Models: Cell Lines |  |  |
| THP-1 | ATCC | TIB-202 |
| MOLT-4 | ATCC | CRL-1582 |
| CAL-1 | Provided by Miwako Narita<sup>4</sup> | CVCL_5G46 |
| HeLa | ATCC | CRM-CCL-2 |
| HEK293T | ATCC | CRL-3216 |
| U937 | ATCC | CRL-1593.2 |
| Kasumi-1 | ATCC | CRL-2724 |
| LP1 | Provide by Jerome Moreaux | CVCL_0012 |
| RPMI 8226 | Provide by Jerome Moreaux | <u>CCL-155</u> |
| Deposited Data |  |  |
| Single cell RNA-seq data | TCGA data base |  |
| Experimental Models: Organisms/Strains |  |  |
| Mouse: C57BL/6J | The Jackson Laboratory | Stock 000664 |
| FlowJo | Treestar | V10.8.1 |
| 7500Software | Applied Biosystem | V2.3 |
| FiJi-ImaJ | System software | Win64 |
| Prism9 | GraphPad | V9.0 |
| Xenabrowser | University of California Santa Cruz | https://xenabrowser.net/ |

### Chemicals

4µ8c (SML0949), Ceapin A7 (SML2330), Bortezomib (5043140001), GSK2656157 (5046510001), rocaglamide (SML0656), and thapsigargin (T7458, 400nM), tunicamycin (SML1287, 100nM), KIRA6, cycloheximide were purchased from Sigma-Aldrich. Harringtonine fom ABCAM (ab141941).

### Cell lines and culture

THP1, MOLT4, CAL1 and B-EBV cell lines were cultured in RPMI1640 . HELA and HEK293T were growth in DMEM . All growth medium have been supplemented with 10% FCS (Biosera). In addition, CAL-1 were grown in the presence of 10mM Hepes, 1mM of sodium pyruvate, 1X of glutamax, 1X non essential amino acids provide by Gibco. For THP1, MOLT4 CAL1 and B-EBV the percentage of FBS were decrease to 1% 16h prior thapsigargin (Tg) and tunicamycin (Tm) treatments. Adherent cells were first trypsinised then washed with ice cold PBS before nuclei extraction. Whereas suspension cells were directly harvested and washed in ice-cold PBS prior viability staining and nuclei extraction.

### Quantitative PCR

Total RNA was extracted from cells using the RNeasy Mini Kit (QIAGEN). cDNA was synthesized using the Superscript II Reverse Transcriptase (Invitrogen) and quantitative PCR were paerformed with ONEGreen FAST qPCR Premix provide by Ozyme using 10µM of each specific primer on a 7500 Fast RealPCR system (Applied Biosystems). cDNA concentration in each sample was normalized to GAPDH expression. The primers used for gene amplification are depicted in the Primers table.

### Immunoblotting

Cells were lysed in Triton buffer (20 mM Tris, pH 7.6, 10 mM NaCl, 1.5mM MgCl2; 1% Triton) supplemented with Complete Mini Protease Inhibitor Mixture Tablets (Roche), NaF (Ser/Thr and acidic phosphatase inhibitor), Na_3_VO_4_ (Tyr and alkaline phosphatase inhibitor) and MG132 (proteasome inhibitor). Protein quantification was performed using the BCA Protein Assay (Pierce). Around 20 μg of soluble proteins were run in 10% acrylamide gels and for the immunoblot the concentration and time of incubation had to be optimized for each individual antibody. Rabbit antibodies against eIF2α, p-eEF2(Thr56), eEF2, eIF2B, p-IRF3 (ser396), IRF3, p-S6, and PERK were purchased from Cell Signaling (ref 5324, 2331, 2332, 3592, 4947, 4302, 2211, and 3192, respectively). Rabbit antibody against p- eIF2α(S51) was purchased from ABCAM (Ref 32157). Rabbit antibody against ATF4 was purchased from Santa Cruz Biotechnology (sc-200). Mouse antibody against β-actin was purchased from Sigma-Aldrich (A2228). Mouse antibodies against HDAC1 and S6 were purchased from Cell Signaling (ref 5356, 2317, respectively). Mouse antibody against puromycin was purchased from Merck Millipore (MABE343). Mouse antibody against p-eIF2β was a kind gift from David Litchfield (University of Western Ontario). Mouse antibody against eIF2β was purchased from Santa Cruz Biotechnology (sc-9978). HRP secondary antibodies were from Jackson ImmunoResearch Laboratories.

### Immunofluorescence confocal microscopy

For immunofluorescence confocal microscopy, cells were seeded on coverslips, fixed with 3.3% PFA and permeabilized 5min with 0.1 % Triton X-100 or 0.05% Saponin. Before staining, samples were incubated with blocking buffer (PBS 1X, 5% FCS, 1% Glycine). Antibodies were added on samples in a wet chamber for 1h at RT or overnight at 4°C. Coverslips were washed in PBS three times before secondary staining. Samples were then washed in PBS and pure water prior glass mounting in ProLong™ Glass Antifade Mountant with nucleic stain (Invitrogen P36980).

### Gene signature generation and MM patient stratification

Gene expression data from purified MM cells were obtained from a publicly available cohort of newly-diagnosed MM patients treated with high dose melphalan and autologous hematopoietic stem cell transplantation (UAMS-TT2 cohort (Accession number GSE24080)). We focused on the expression of genes regulated by XBP1, ATF6, and ATF4. We also used Affymetrix data from relapsed MM patients at relapse subsequently treated with bortezomib or dexamethasone monotherapy (Accession number GSE9782) from the study by Mulligan et al ^40^. We also used a cohort of MM patients at relapse treated by anti-CD38 MoAb^55,56^. MM cells were purified using anti-CD138 MACS microbeads (Miltenyi Biotec, Bergisch Gladbach, Germany) and their gene expression profile (GEP) obtained using Affymetrix U133 plus 2.0 microarrays as previously described^56,57^. Gene expression data were normalized with the MAS5 algorithm and analyses processed with GenomicScape (http://www.genomicscape.com)^58^. All statistical analyses were performed with the statistics software R, and R packages developed by Bioconductor project (https://www.bioconductor.org) (ref). The target genes were selected based on their association with these transcription factors as identified through chromatin immunoprecipitation (ChIP) assays and from QIAGEN IPA databases. The XBP1-Pathway signature was computed with 23 XBP1-target genes associated with a prognostic value in MM patients. The XBP1-Pathway signature is defined by the sum of the beta coefficient derived from the Cox model for each prognostic gene weighted by − 1 or + 1 according to the MMC gene expression above or below the Maxstat R package defined cutpoint as previous ly described^59,60^. The statistical significance of differences in overall survival or progression free survival between patients’ groups was calculated using the log-rank test and survival curves were plotted using the Kaplan-Meier method.

### Cytotoxicity assays

A total of 20,000 cells per well were seeded in 96-well plates using complete RPMI medium supplemented with 10% fetal calf serum (FCS). Cells were treated in triplicate with varying concentrations of bortezomib (catalog number 5043140001), 4µ8c (catalog number SML0949), Ceapin A7 (catalog number SML2330), and GSK2656157 (catalog number 5046510001). The treatments were administered to evaluate the combined cytotoxic effects under different drug concentration conditions. Cell viability was assessed at 24 and 48 hours post-treatment. A supra-vital viability dye, Zombie Yellow™ (1:200 dilution, catalog number 423103), was employed to stain dead cells. Following staining, the cells were washed thoroughly to remove excess dye and then subjected to flow cytometric analysis.

### Statistical analysis

All statistics were done using Prism 9 software. The most appropriate statistical test was chosen according to each set of data, as indicated in f<u>igure</u> legends with p values *p<0.05; **p<0.01; ***p<0.001; ****p<0.0001.

## Acknowledgments

We thank Dr. Pierre Golstein and Dr. Ludovic Martinet for critical reading of the manuscript. We want to thank Sylvain Bigot and Cyrille Mionnet from the CIML cytometry platform.

## Funding

This research was supported by CENTURI postdoc funding P.G.G Supported by the ANR JCJC-Epic ZENITH (RJA)

ANR-20-CE14-0028 to RJA

ANR PRC MetaNiche ANR-22-CE15-0015-02 to RJA

European Commission Horizon2020 Transcan 2021-227-TALETE French National Cancer Institute (INCa) to RJA.

This work was partially funded by the French National Cancer Institute (INCa) SIRIC – Pediacriex “SOUTH-Rock”.

## Author Contributions

Conceptualization: R.J.A., J.P.G.;

Supervision: R.J.A and P.P.;

Resources: J.M., S.R., M.N,

Investigation: J.P.G, P.G.G, L.G., A.R., F.F.S, E.S., D.B.D.S., S.F, Y.G.

Bioinformatics and Figures: J.P.G, A.C, R.J.A, J.M., P.P.

Writing – original draft: R.J.A, J.P.G.

Writing review & editing: P.P, P.G.G, F.F.S, R.L.H, B.N., E.G., A.C.

Funding acquisition: RJA and P.P.

## DECLARATION OF INTERESTS

The authors declare no competing interests.

## SUPPLEMENTAL FIGURES

**Supplemental Figure 1.**
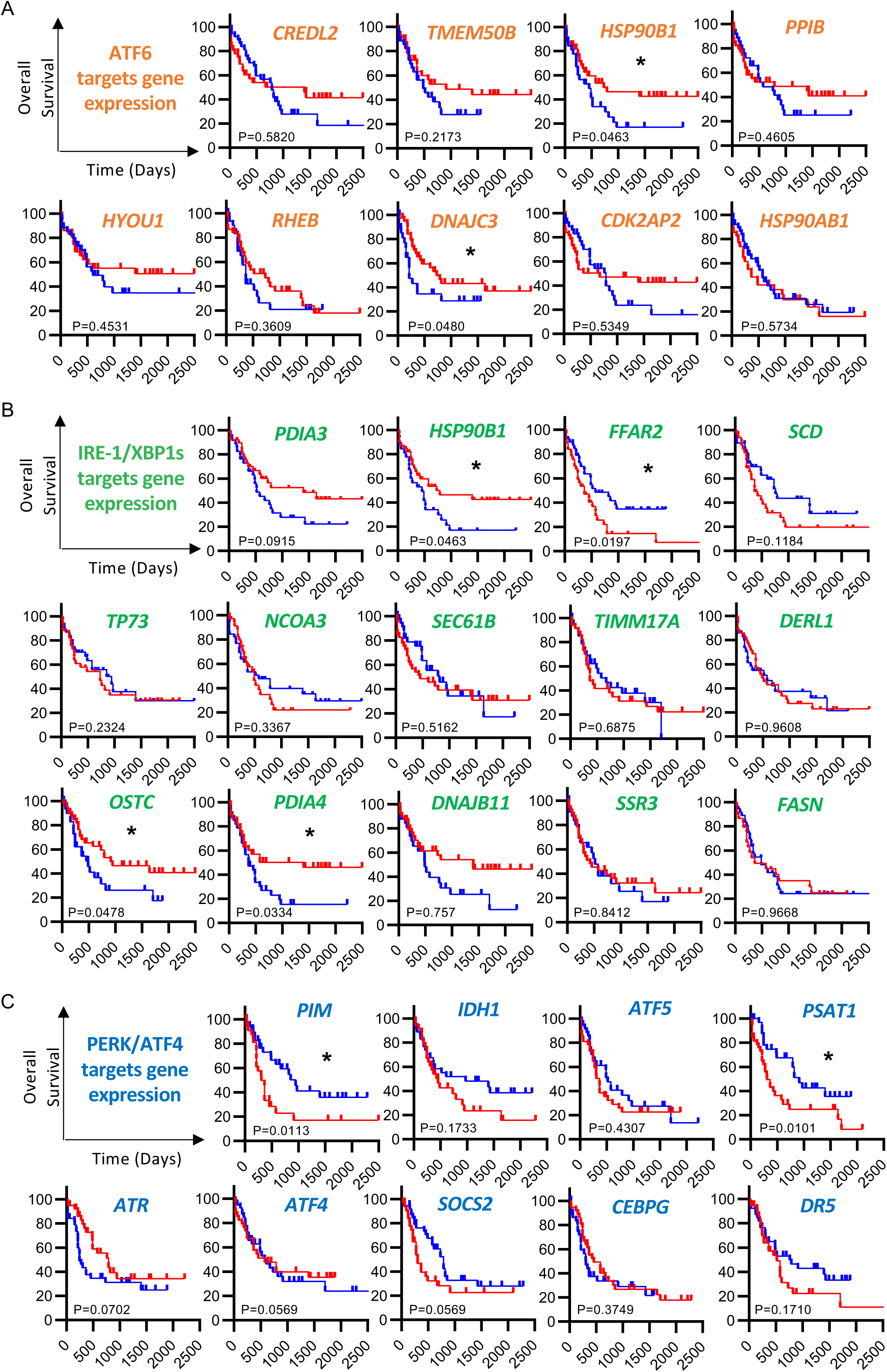
Branch-specific UPR signatures correlate with survival prognosis of patients with Acute Myeloid Leukemia. (A-C) Kaplan Meier survival curves for genes under the control of ATF6, XBP1s or PERK generated for AML. Kaplan Meier curves were done based on dichotomized gene expression, specifically for values below quartile 1 (blue) and above quartile 3 (red). Genes indicated are described to be under the control of ATF6 (A); IRE- 1/XBP1s (B) and PERK/ATF4 (C). Statistical analysis was performed using Log-rank test. \**P* < 0.05.

**Supplemental Figure 2.**
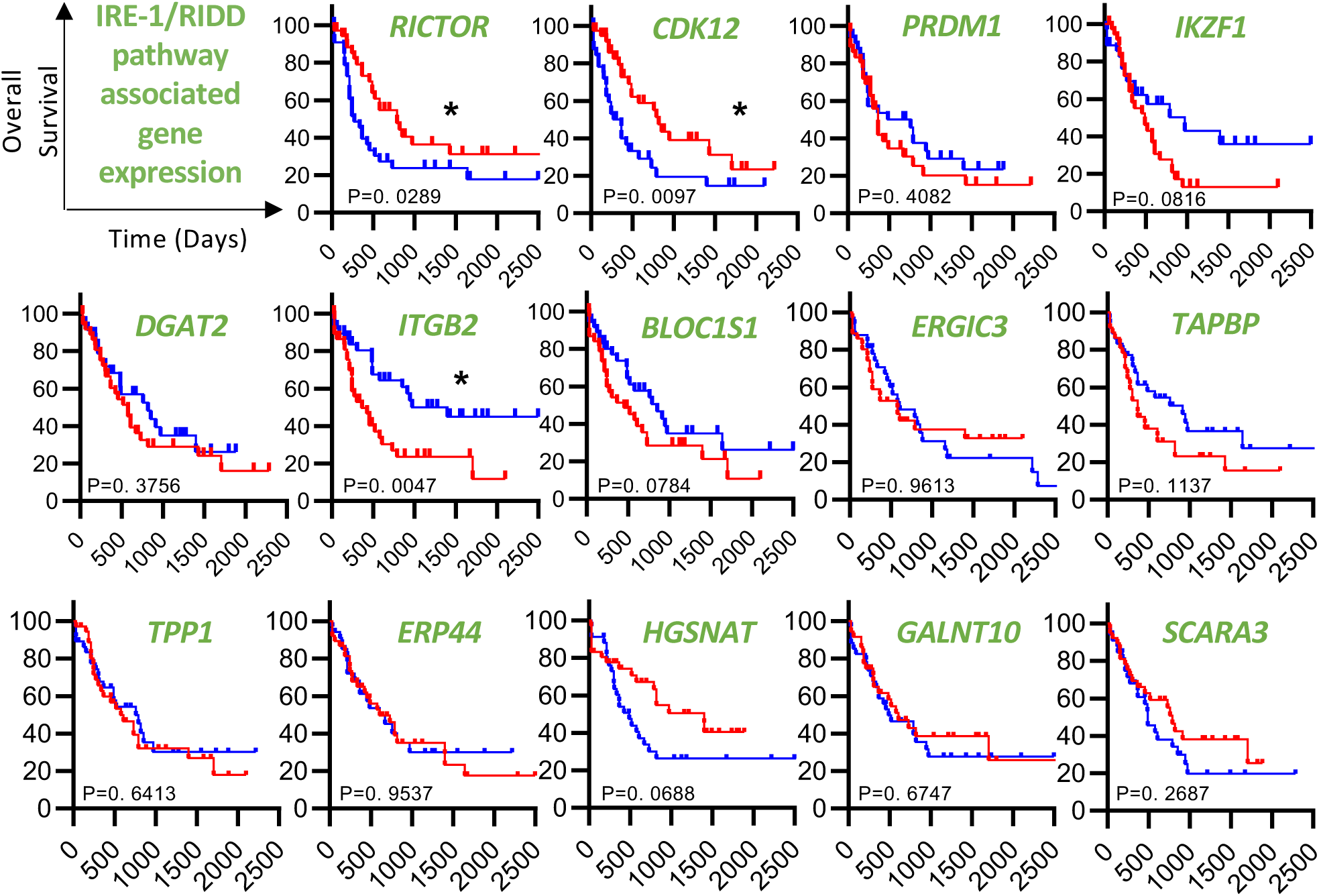
RIDD targets expression correlate with survival prognosis of AML patients. Kaplan Meier survival curves for mRNAs targeted by RIDD were generated for AML. Kaplan Meier curves were done based on dichotomized gene expression, specifically for values above quartile 1 (blue) and quartile 3 (red). Statistical analysis was performed using Log-rank test. \**P* < 0.05.

**Supplemental Figure 3.**
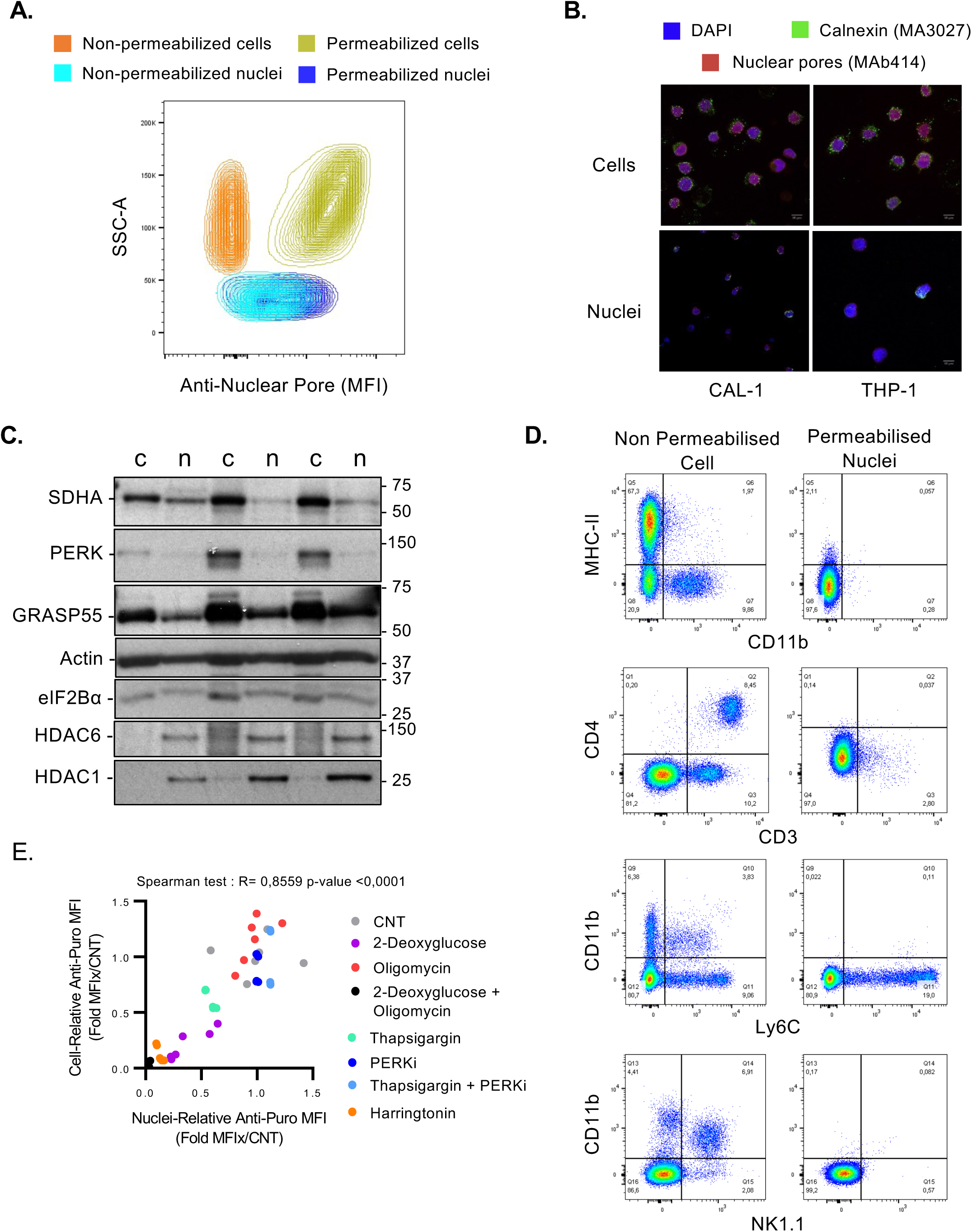
Quality control of nuclear extraction for SNUPR profiling. **(A)** Nuclei or whole CAL-1 cells were permeabilized or not and stained with nucleopore antibody. (**B**) Staining of nuclear pores and calnexin was analysed by microscopy in CAL-1 and THP-1 whole cells and nuclei. (**C**) CAL-1 cells (c) or nuclei (n) suspensions were lysed and analysed by immunoblot. Relative quantification of each protein is indicated on the right side of the panel. (**D**) Whole cells and nuclei extract from mouse blood stained with indicated surface markers. (**E**) Correlation of puromycin MFI (normalized to control) of cells versus isolated nuclei in response to different pharmacological compounds. Spearman test: R= 0,8559 p-value <0,0001. CNT: Control.

**Supplemental Figure 4.**
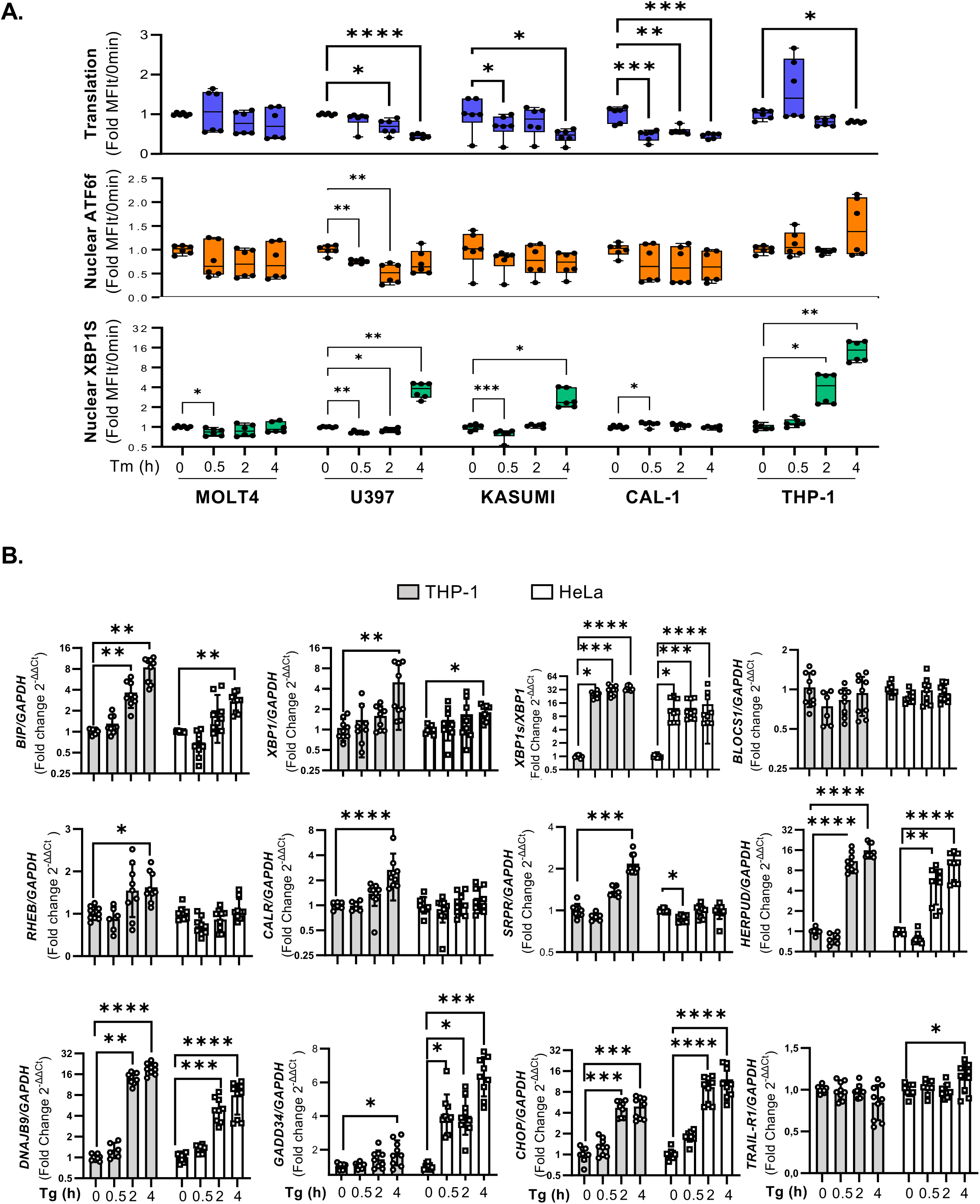
R**a**pid **and efficient inhibition of protein synthesis modulates UPR activation. (A)** Distinct cancer cell lines were treated with 100ng/ml of tunicamycin (Tm) for 30min, 2h and 4h prior to nuclei extraction and further SNUPR profiling of UPR activation by measuring translation (puromycin incorporation) and translocation of ATF6f and XBP1s by flow cytometry. Statistical analysis was performed using Kruskall-Wallis test for each cell line. \**P* < 0.05, \*\**P* < 0.01, \*\*\**P* < 0.001 and \*\*\*\**P* < 0.0001. **(B)** qPCR quantification of relative mRNA levels of UPR-associated transcripts on HeLa and THP-1 cells treated for 30min, 2h and 4h with thapsigargin (Tg, 400nM).

**Supplemental Figure 5.**
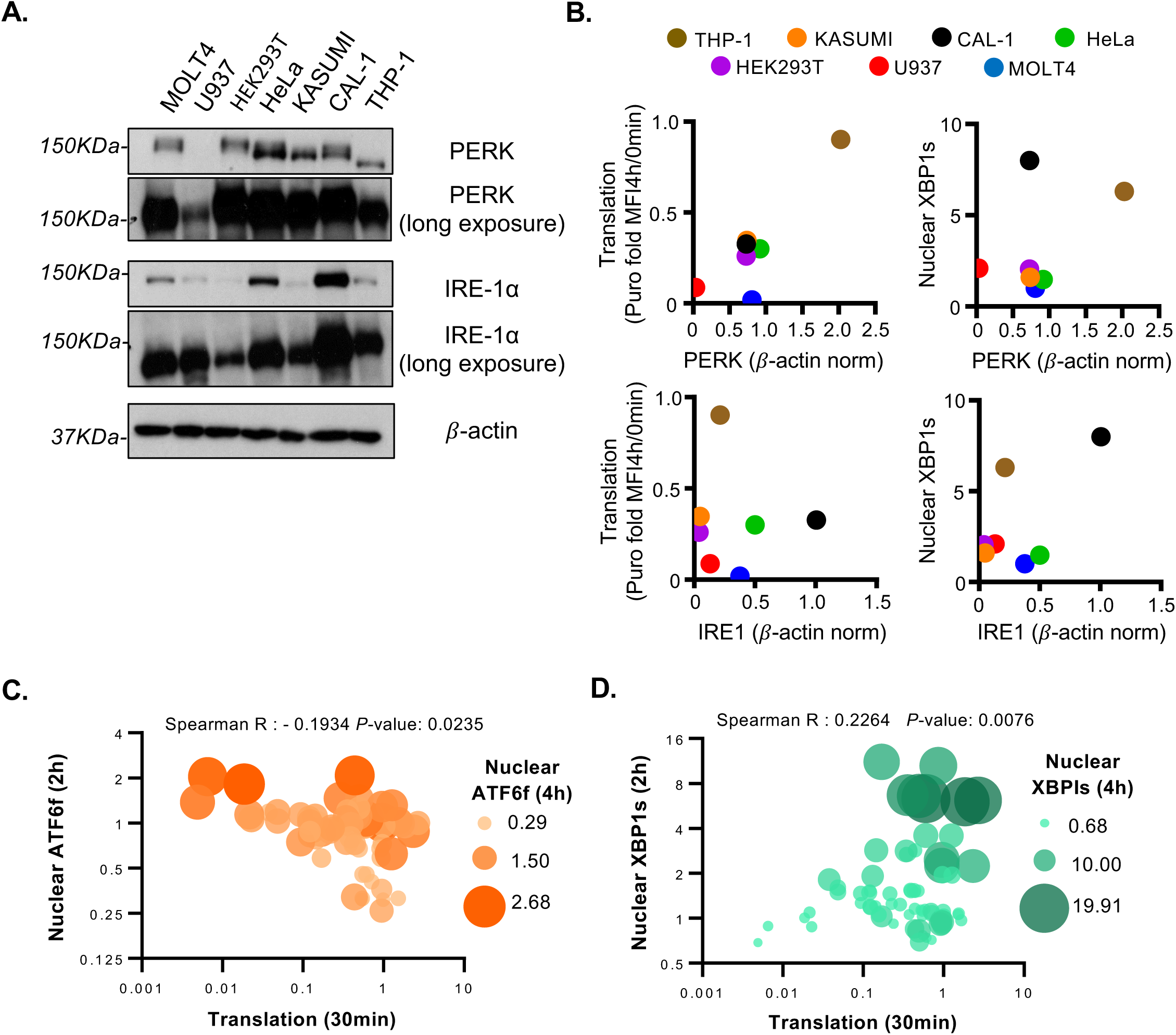
C**o**rrelation **of translation inhibition with reduced XBP1s expression. (A)** Quantification of IRE-1a and PERK protein levels on different cell lines was assessed by immunoblot. **(B)** Correlation analyses between expression levels of IRE-1 and PERK with translation levels or nuclear translocation of XBP1s for each cell line was assessed. Further correlation analyses between early translation levels (30min) and nuclear expression of **(C)** ATF6 and **(D)** XBP1s after 2h of ER stress induction was evaluated using the Spearman test. Spearman R-score and p-values are depicted in the lower panel.

**Supplemental Figure 6.**
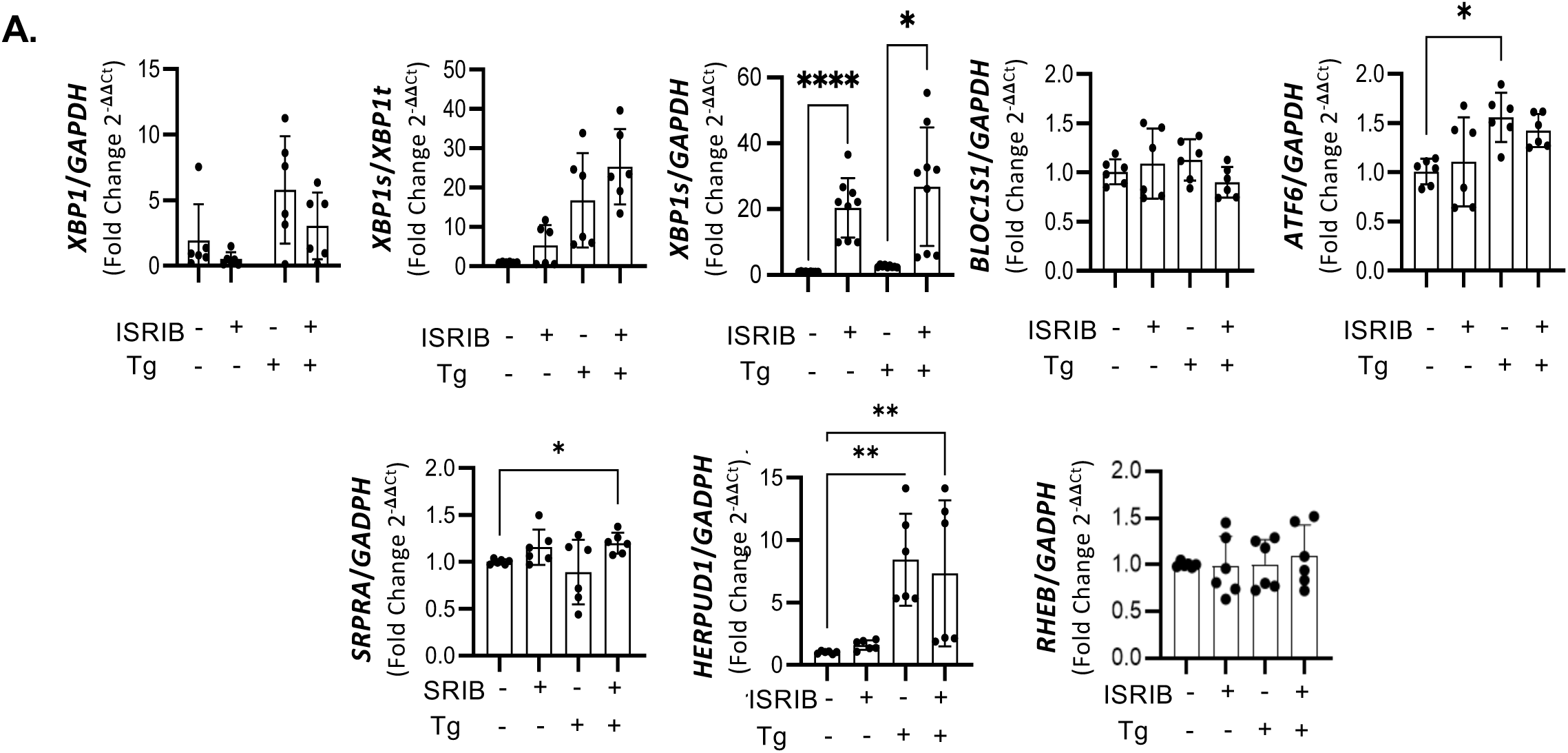
T**r**anslation **inhibition delays IRE-1/XBP1 axis activation.** HeLa cells were treated 30min or 4h with thapsigargin (Tg, 400nM) in the presence or absence of ISRIB (1ug/mL) prior to nuclei and RNA isolation for SNUPR profiling and transcriptional analysis of UPR-associated transcripts. **(A)** RT-qPCR analysis of XBP1 splicing as well as relative mRNA levels of XBP1s, XBP1tot, ATF6a and BLOC1S1 on HeLa cells treated with Tg in combination with ISRIB. Statistical analysis was performed using 2way ANNOVA test. \**P* < 0.05, \*\**P* < 0.01, and \*\*\**P* < 0.001.

**Supplemental Figure 7.**
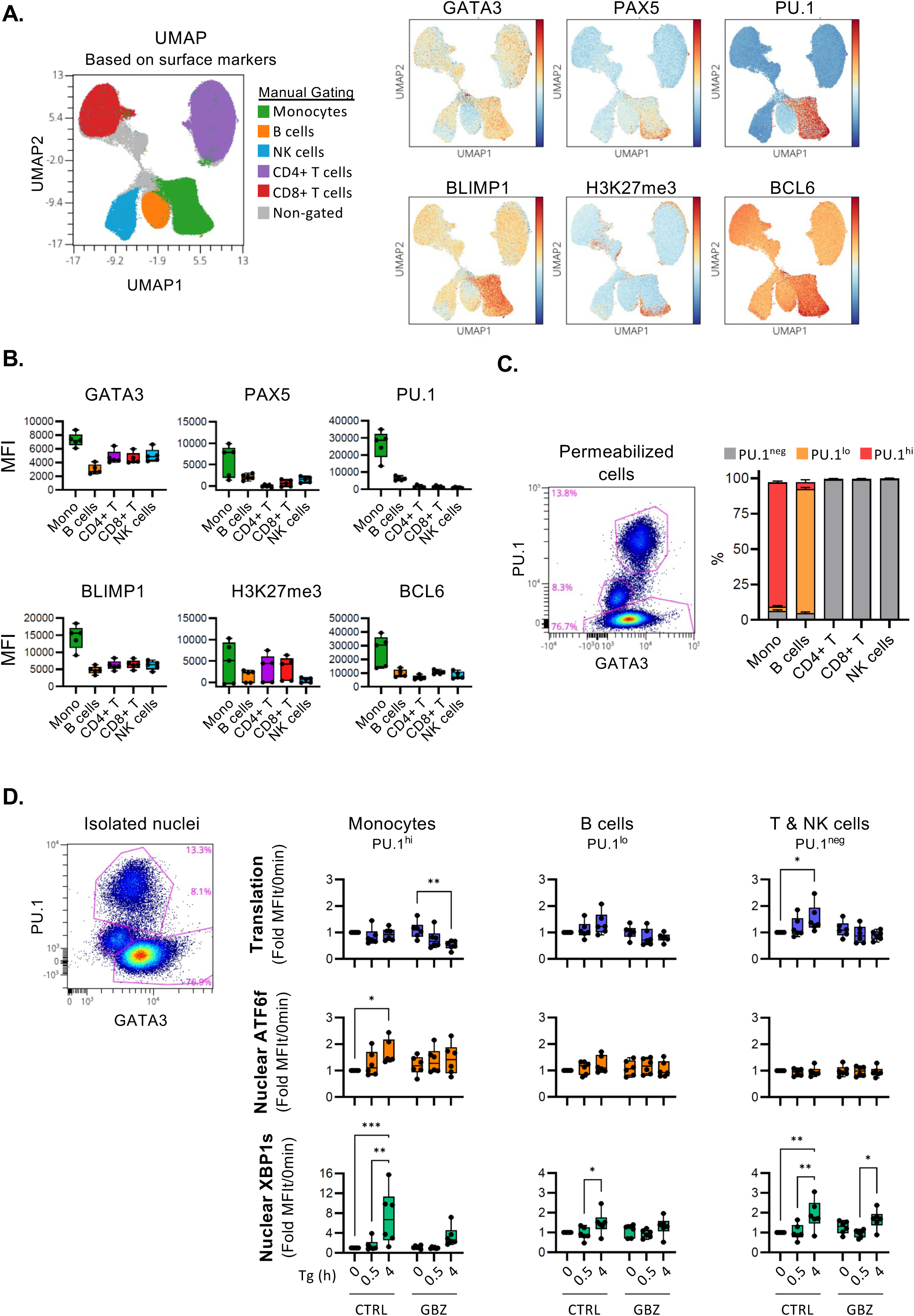
L**i**neage**-associated transcription factor staining allows SNUPR profiling on specific cell subsets on PBMC.** Human Peripheral Blood Mononuclear Cells (PBMC) from 6 healthy donors were stained with surface markers (CD3, CD4, CD8, CD19, CD16, HLA-DR), permeabilized and stained intracellularly for nuclear markers (GATA3, PAX5, PU.1, BLIMP1, H3K27me3, BCL6) to identify cell subsets by FACS. **(A)** UMAP for PBMC was generated by using surface markers only. UMAPs colored either by manually gated immune populations based on surface markers (left); or by nuclear marker expression levels (right panel). **(B)** Quantification of MFI for different nuclear markers on main immune populations. **(C)** Three gates were defined based on PU.1 and GATA3 expression levels for permeabilized cells. Stacked bars show the distribution of cells in these gates for each immune cell subset. **(D)** SNUPR profiling of PBMC. PBMCs were treated with thapsigargin (Tg, 400nM) for 30min or 4h in the presence or absence of guanabenz (GBZ, 50µM) prior to nuclear isolation. Nuclei identity was defined based on PU.1 and GATA3 expression (left). Translation levels and nuclear translocation of ATF6f and XBP1s was measured on isolated nuclei by FACS (right). MFI values were normalized to control treated samples. Statistical analysis was performed using 2way ANNOVA test. *P < 0.05, **P < 0.01, and ***P < 0.001.

**Supplemental Figure 8.**
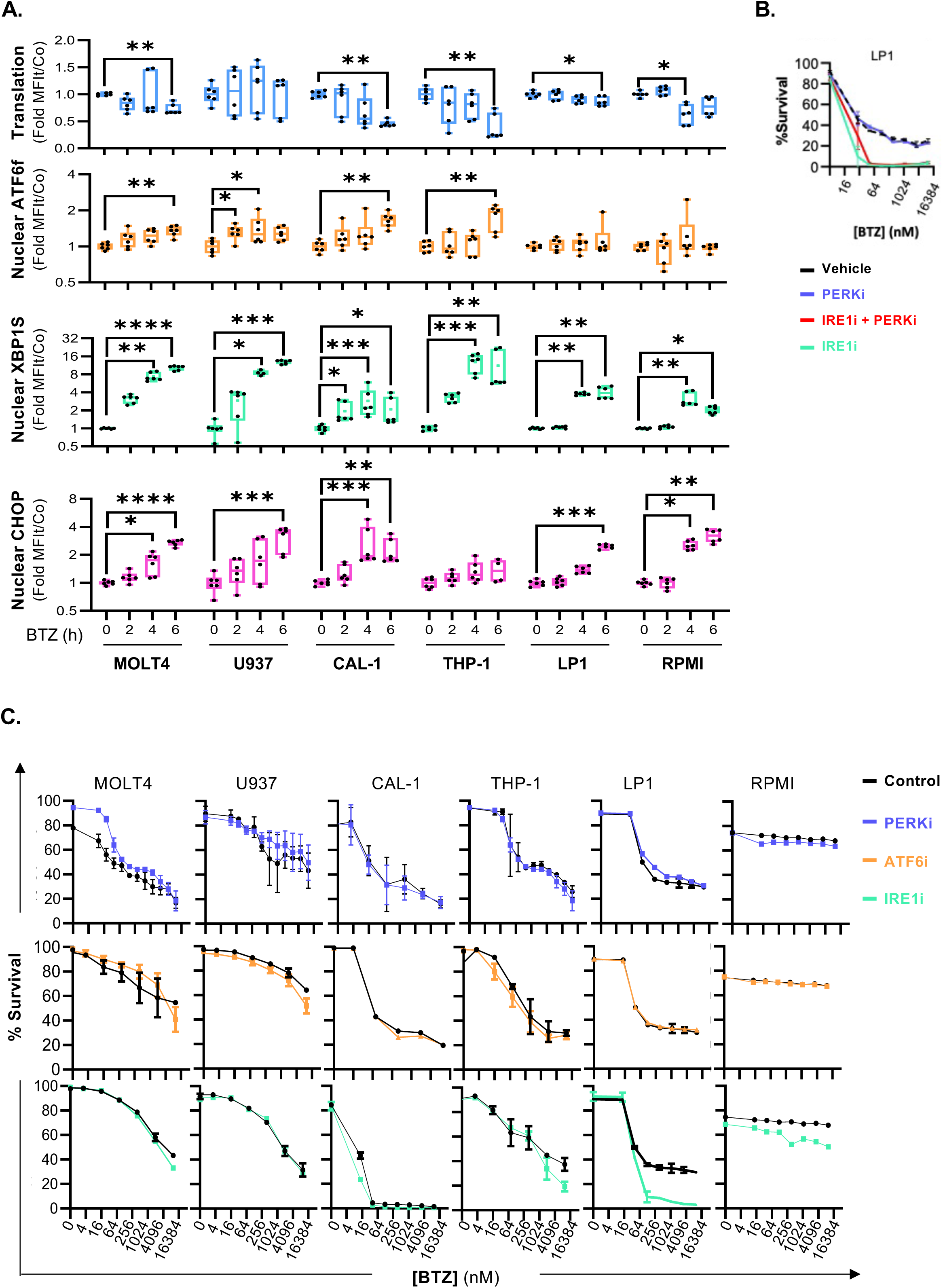
B**o**rtezomib **induces UPR activation in leukemic and myeloma cells.** Leukaemia cell lines and two multiple myeloma (MM) cell lines, LP1 and RPMI, were treated with the proteasome inhibitor bortezomib (BTZ, 100nM) for 2h, 4h and 6h prior to nuclei isolation and SNUPR profiling. **(A)** SNUPR profiles of translation levels and ATF6f and XBP1s nuclear translocation of cancer cells treated with BTZ for 2h, 4h or 6h. Statistical analysis was performed using Kruskall-Wallis test for each cell lines. \**P* < 0.05, \*\**P* < 0.01, and \*\*\**P* < 0.001. **(B)** Cell survival analysis of LP1 cells treated 6h with BTZ in presence or absence of IRE-1 inhibitor 4u8c (10uM). PERK inhibitor GSK2656157 (100nM) or a combination of both measured by flow cytometry. **(C)** Survival analysis of leukemia and multiple myeloma cells treated with a gradient concentration of BTZ in combination with a PERK inhibitor (GSK2656157. 100nM. blue). ATF6 inhibitor (Ceapin A7, 6uM orange) or IRE-1 inhibitor (4u8, green) for 24h. All analysis were performed by flow cytometry measurements of cell viability dye signals.

**Supplemental Figure 9.**
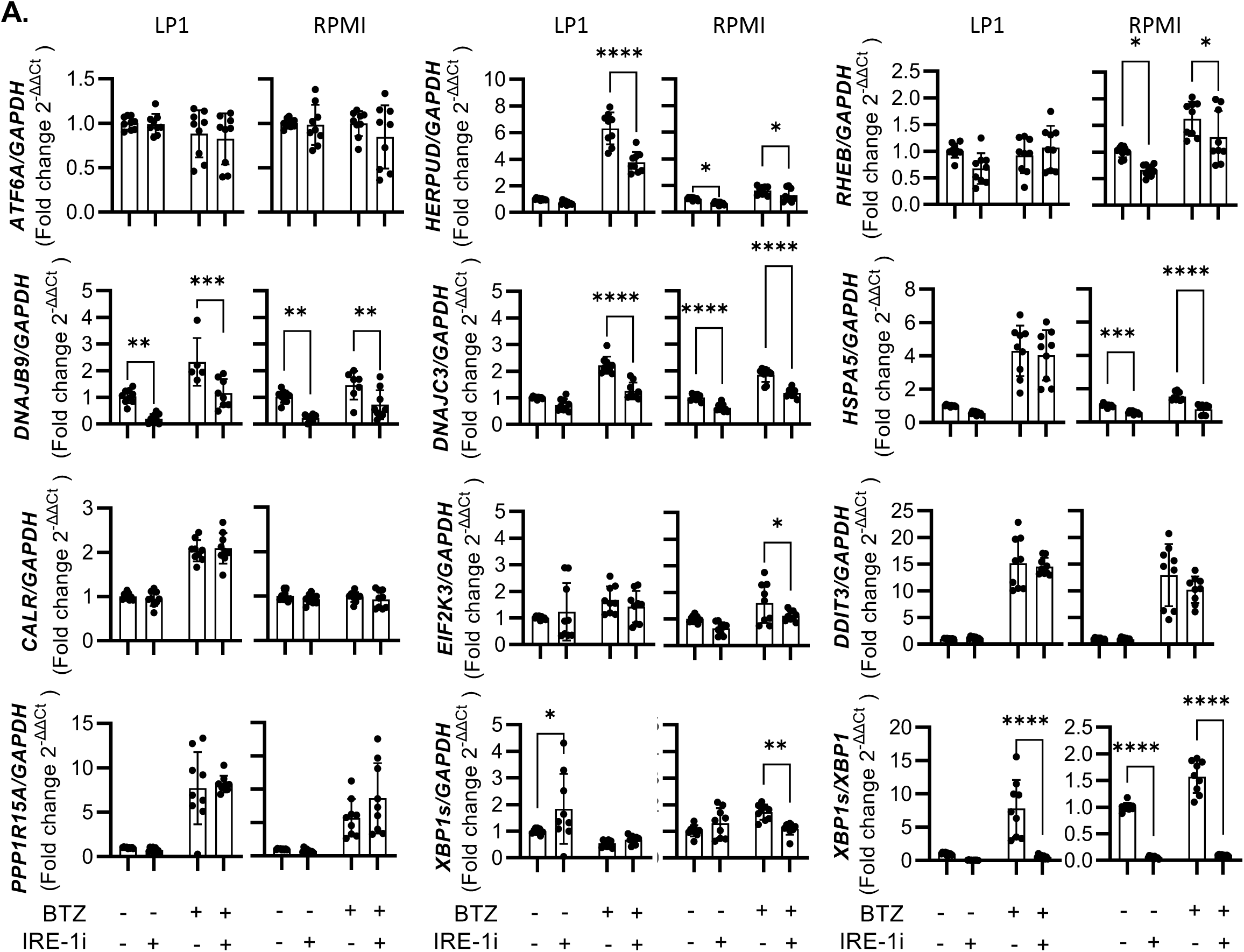
I**R**E**-1 inhibition affects ER chaperon expression on MM cells. (A)** RT-qPCR analysis of ER chaperones and other UPR-associated transcript mRNA levels of LP1 and RPMI MM cell lines treated with BTZ (100nM) for 4h in the presence or absence of IRE-1 inhibitor 4µ8c (10µM). Statistical analysis was performed using 2way Annova test. \**P* < 0.05, \*\**P* < 0.01, and \*\*\**P* < 0.001.

**Supplemental Table 2.**
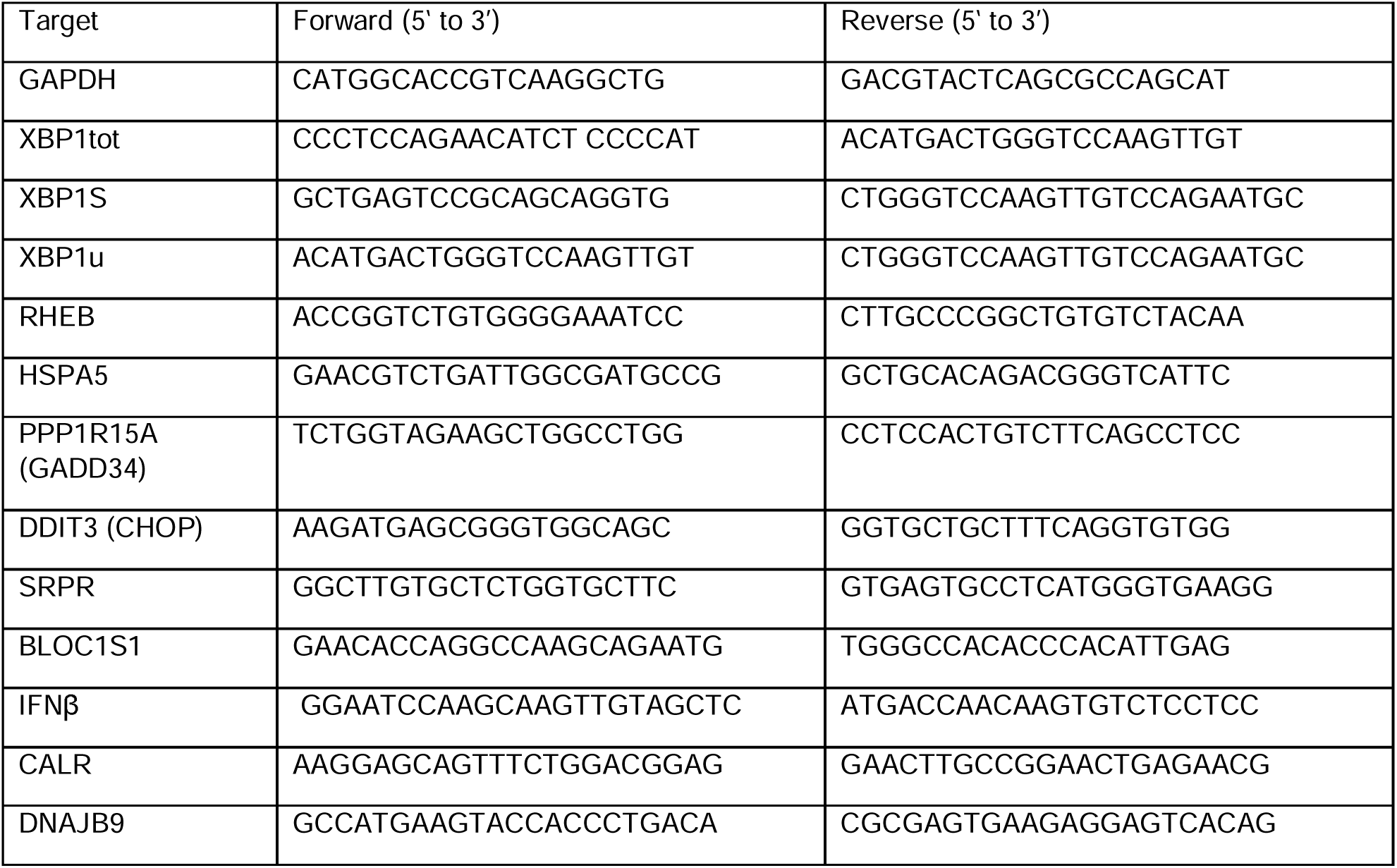
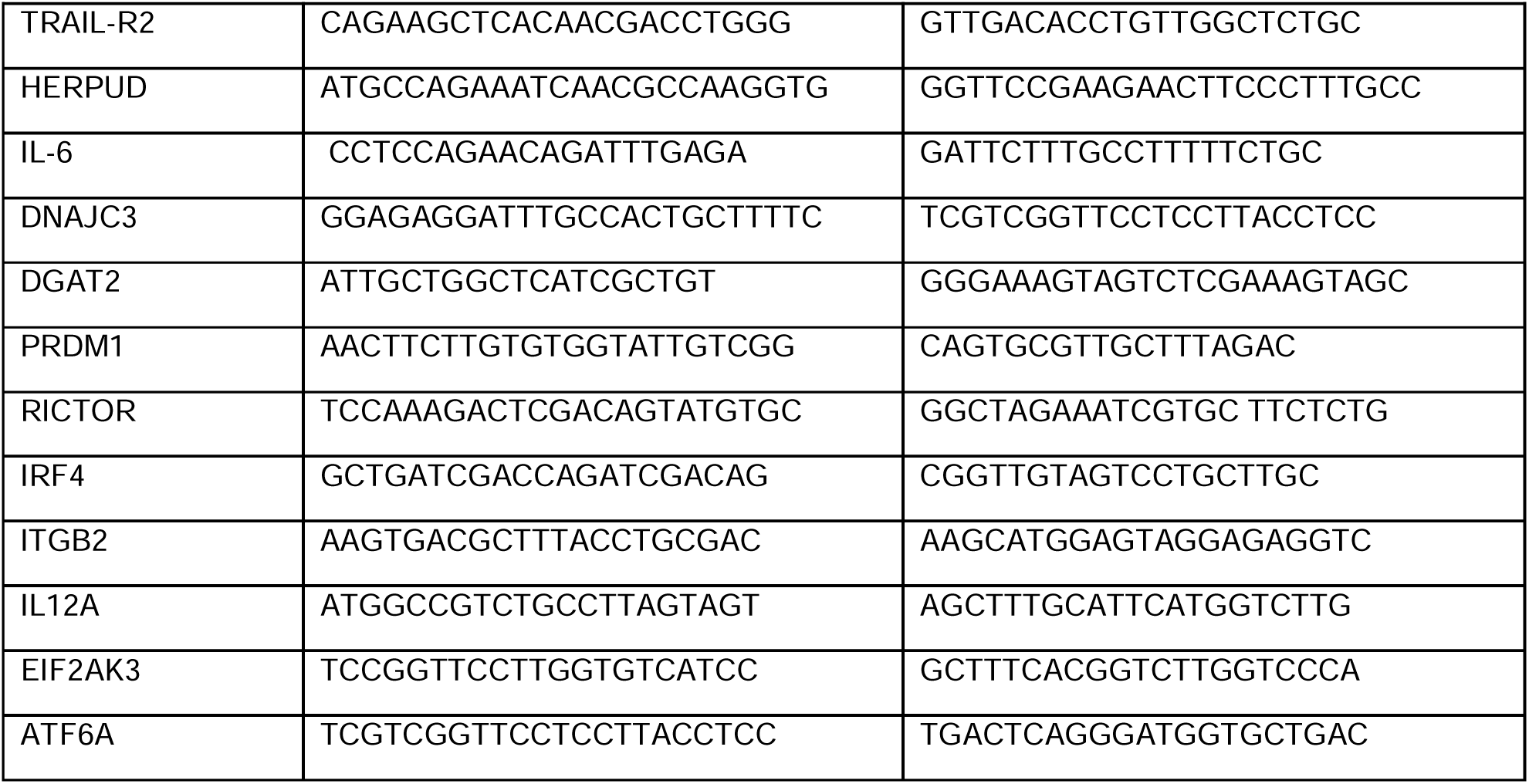
Primers used.

## ACKNOWLEDGEMENTS

Revision of Manuscript and comments: Dr. Pierre Golstein and Dr. Ludovic Martinet.

## RESOURCE AVAILABILITY

### Lead contact

Further information and requests for resources and reagents should be directed to and will be fulfilled by the lead contact. Rafael J. ARGÜELLO

### Materials availability

Materials generated in this study will be made available on request, but we may require a completed Materials Transfer Agreement.

### Data and code availability

Original western blot, gel, and microscopy images have been deposited at Mendeley and are publicly available as of the date of publication. The DOI is listed in the key resources table.

This paper does not report original code.

Any additional information required to re-analyze the data reported in this paper is available from the lead contact upon request.

## Notes

### Competing Interest Statement

The authors have declared no competing interest.

### Summary of Updates

We modified the order of authors according to updated contributions.

